# Proteome-Scale Relationships Between Local Amino Acid Composition and Protein Fates and Functions

**DOI:** 10.1101/338202

**Authors:** Sean M. Cascarina, Eric D. Ross

## Abstract

Proteins with low-complexity domains continue to emerge as key players in both normal and pathological cellular processes. Although low-complexity domains are often grouped into a single class, individual low-complexity domains can differ substantially with respect to amino acid composition. These differences may strongly influence the physical properties, cellular regulation, and molecular functions of low-complexity domains. Therefore, we developed a bioinformatic approach to explore relationships between amino acid composition, protein metabolism, and protein function. We find that local compositional enrichment within protein sequences affects the translation efficiency, abundance, half-life, subcellular localization, and molecular functions of proteins on a proteome-wide scale. However, these effects depend upon the type of amino acid enriched in a given sequence, highlighting the importance of distinguishing between different types of low-complexity domains. Furthermore, many of these effects are discernible at amino acid compositions below those required for classification as low-complexity or statistically-biased by traditional methods and in the absence of homopolymeric amino acid repeats, indicating that thresholds employed by classical methods may not reflect biologically relevant criteria. Application of our analyses to composition-driven processes, such as the formation of membraneless organelles, reveals distinct composition profiles even for closely related organelles. Collectively, these results provide a unique perspective and detailed insights into relationships between amino acid composition, protein metabolism, and protein functions.

**Author Summary:** Low-complexity domains in protein sequences are regions that are composed of only a few amino acids in the protein “alphabet”. These domains often have unique chemical properties and play important biological roles in both normal and disease-related processes.While a number of approaches have been developed to define low-complexity domains, these methods each possess conceptual limitations. Therefore, we developed a complementary approach that focuses on local amino acid composition (i.e. the amino acid composition within small regions of proteins). We find that high local composition of individual amino acids is associated with pervasive effects on protein metabolism, subcellular localization, and molecular function on a proteome-wide scale. Importantly, the nature of the effects depend on the type of amino acid enriched within the examined domains, and are observable in the absence of classically-defined low-complexity (and related) domains. Furthermore, we define the compositions of proteins involved in the formation of membraneless, protein-rich organelles such as stress granules and P-bodies. Our results provide a coherent view and unprecedented resolution of the effects of local amino acid enrichment on protein biology.

## Introduction

Low-complexity domains (LCDs) in proteins are regions enriched in only a subset of possible amino acids. LCDs can be composed of homopolymeric repeats of a single amino acid, short tandem repeats consisting of only a few different amino acids, or aperiodic stretches with little amino acid diversity [1]. Proteins containing LCDs are relatively common among organisms from all domains of life, and are particularly common among eukaryotes [2–4]. For example, approximately 70% of genes in the *Saccharomyces cerevisiae* genome possess at least one classically-defined LCD [3]. Furthermore, the total number of LCDs far exceed the total number of yeast genes (∼2-fold more LCDs than genes), indicating that many genes contain multiple distinct LCDs.

Various methods have been developed to assess biopolymer sequence complexity [1,5–9]. One of the most commonly employed methods to define LCDs is the SEG algorithm [1], which scans protein (or nucleic acid) sequences using a short sliding window, and calculates the local Shannon entropy for each window (see [10] for a detailed description). Subsequences with a Shannon entropy value below a pre-determined “trigger” threshold are classified as LCDs. LCD boundaries are later extended and refined by merging overlapping LCDs and calculating combinatorial sequence probabilities. Another metric commonly used to assess relative sequence complexity is compositional bias, which involves determining the statistical probability of a sequence given whole-proteome frequencies of the individual amino acids [11,12]. These approaches (or closely-related approaches) have been used extensively to examine LCDs on a proteome-wide scale [1,3,12–17].

LCD-containing proteins have been implicated in a variety of normal and pathological cellular processes. For example, Q/N-rich yeast proteins often play a role in transcription regulation, endocytosis, and cell cycle regulation, among other functions [11,18]. Many proteins containing Q/N-rich LCDs, or LCDs of related types (Q/N/G/S/Y-rich LCDs) have been linked to prion or prion-related processes [11,18–21]. Additionally, many prion-like LCDs have been linked to stress granules and processing bodies (P-bodies) in eukaryotes (see [22] for recent review). The amino acid composition of these LCDs confers unusual biophysical properties to these domains [23], which likely relates to their unique behavior *in vitro* and *in vivo* [24–29].However, these unusual characteristics appear to be inseparably linked to pathological processes as well. For example, genetic expansion of regions encoding homopolymeric glutamine repeats (the simplest type of LCD) in various proteins can lead to a multitude of neurodegenerative disorders, including Huntington’s Disease and spinocerebellar ataxias (for review, see Weber et al., 2014). Furthermore, mutations in the LCDs of stress granule proteins can alter stress granule dynamics and lead to degenerative diseases [25,27,29,31,32]. The importance of LCDs extends well beyond Q/N-rich LCDs, as LCDs of other compositions have also been linked to normal and pathological cellular processes [12,14,17,33,34].

Although LCDs can clearly impact protein regulation and function, a number of challenges have thus far limited a proteome-scale understanding of these relationships. One major challenge lies in defining LCDs. Current approaches use statistically-defined thresholds for sequence complexity or compositional bias [1,11], or arbitrarily-chosen repeat lengths for proteins with homopolymeric repeats [33–39]. Although these definitions of LCDs, compositionally biased sequences (herein referred to as “statistically-biased domains” to avoid later confusion), or homopolymeric repeats have facilitated important discoveries, the biological relevance of these thresholds has not been rigorously examined. Furthermore, these proteins are often grouped into a single class even though their compositions, and therefore physical properties, can differ dramatically (a limitation that was appreciated in a recent review [40]).

To address these limitations, we have developed an alternative approach to infer relationships between amino acid composition and protein metabolism and function. By focusing on amino acid composition, which is the fundamental feature underlying both sequence complexity and statistical amino acid bias, we examined the effects of local compositional enrichment on various aspects of protein regulation and function without appealing to pre-defined sequence complexity or statistical bias thresholds. We find that local compositional enrichment affects nearly all core aspects of a protein’s tenure in the cell, including translation efficiency, abundance, half-life, subcellular localization, and function. However, enrichment for different amino acids leads to different effects, even for residues often grouped based on physicochemical similarities, highlighting the importance of distinguishing LCDs of different types. These effects are discernible at compositions below those required for classification as low-complexity or statistically-biased, suggesting that the thresholds in traditional methods may not be biologically optimized. Finally, analysis of experimentally-defined protein components of stress granules and P-bodies reveals both shared and distinct compositional features associated with these organelles.

## Results

### Systematic Survey of Local Amino Acid Composition

Fundamentally, both sequence complexity and statistical amino acid bias are indirect measures of local amino acid composition. Since composition is a more direct indication of overall protein domain properties, we sought to examine whether composition alone could be used to infer residue-specific relationships between local amino acid composition and protein regulation and function. We first developed an algorithm to partition the yeast proteome on the basis of maximum local composition for each amino acid using a series of scanning window sizes (Fig 1; see Methods). For all amino acids, the majority of proteins are partitioned into composition bins of ≤ 25% (Fig 2 and Table S1). However, the number of proteins achieving higher local compositions, indicated by a right-hand shoulder or tail in the distribution, were strongly residue-dependent. For example, proteins containing local enrichment of highly hydrophobic residues (I, L, M, and V), aromatic residues (F, W, and Y), or cysteine are almost exclusively limited to composition bins of ≤ 45% for the smallest window size, whereas alanine and proline distributions extend to slightly higher composition ranges (up to 60-65%). Proteins containing local enrichment of polar (G, N, Q, S, and T) or charged (D, E, and K) residues in composition bins of ≥ 40% are relatively common even among larger window sizes (albeit to differing degrees), whereas histidine and arginine rich regions are relatively rare. These data indicate that relatively high local enrichment is tolerated for some amino acids, while compositional enrichment for other amino acids appears to be restricted in yeast.

**Fig 1.**
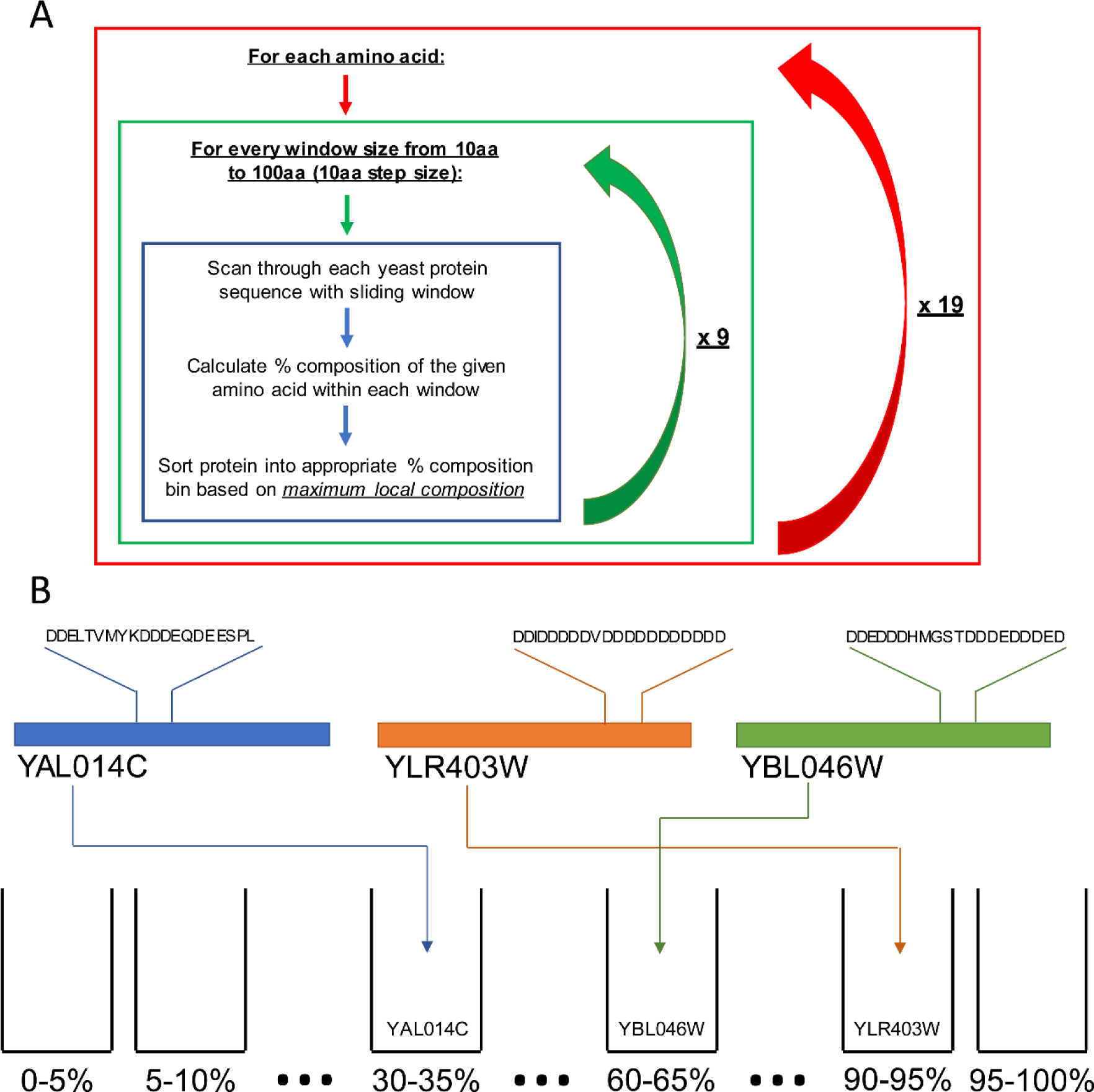
Depiction of proteome sorting on the basis of maximum local composition. (A) For each amino acid and window size combination, each yeast protein is sorted into percent composition bins based on the maximum local composition of the amino acid within the given sliding window size. This effectively sorts the yeast proteome 200 distinct ways (20 amino acids x 10 different sliding window sizes). (B) Visual representation of proteins sorted based on maximum local aspartic acid composition with a 20 amino acid sliding window.

**Fig 2.**
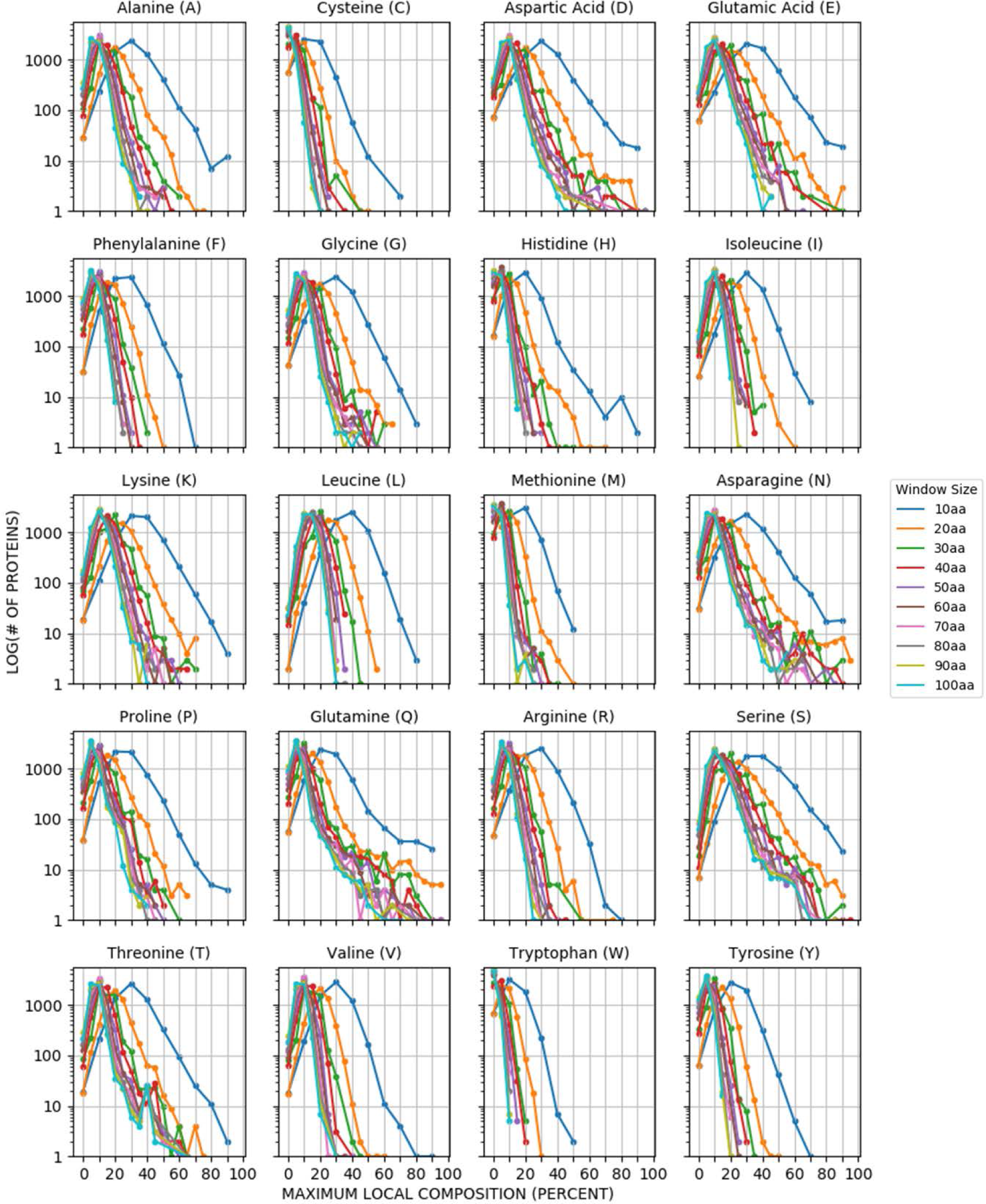
Distribution of the yeast proteome based on maximum local amino acid composition. Protein counts indicate the log of the number of proteins partitioned into each window size/percent composition bin for each of the 20 canonical amino acids. Scatter points are connected by line segments for visual clarity only.

### Local Compositional Enrichment Affects Protein Metabolism in a Residue-Specific Manner

While the origins and evolution of LCDs have been extensively explored [3,4,14,37,41], the regulation and metabolism of LCD-containing proteins remain poorly-understood. Proteins with intrinsically disordered segments, which often qualify as LCDs, have been associated with lower protein half-lives [42]. However, not all intrinsically disordered regions lead to short protein half-lives, and not all LCDs are intrinsically disordered [15]. Additionally, proteins with homopolymeric repeats, when considered as a single class, are associated with lower translation efficiency, lower protein abundance, and lower protein half-life compared to proteins lacking homopolymeric repeats [36]. However, the regulation and structural properties of proteins with LCDs or homopolymeric repeats is likely strongly dependent on the predominant amino acids within the domain of interest [40].

To explore relationships between local compositional enrichment and protein metabolism, we first examined the effects of local compositional enrichment on protein abundance and turnover. Recent advances in proteomic methods have facilitated remarkable proteome coverage for both protein abundance [43] and protein half-life [44] measurements in yeast. At each window size/percent composition bin, the protein abundance distribution of all proteins partitioned into that bin was compared to all other yeast proteins using a Mann-Whitney *U* test. Transitions from significantly lower median abundance to significantly higher median abundance or vice versa are observed upon enrichment for many amino acids individually (Fig 3). However, the effects of progressive compositional enrichment are dependent on amino acid type. For the majority of amino acids (C, D, F, H, I, L, M, N, P, Q, R, S, T, W, or Y) compositional enrichment is associated with lower median protein abundance. However, compositional enrichment of A, G, or V is associated with higher median protein abundance. Two very similar transitions are observed for both E-rich and K-rich sequences: as compositional enrichment increases, the relative median protein abundance transitions from high to low, then back to high. Collectively, these trends are consistent with, yet much stronger than, previously observed correlations between protein abundance and whole-protein composition [45,46], suggesting that the trends observed previously may actually reflect the effects of local compositional enrichment, which would increase apparent whole-protein composition for the enriched amino acid yet be dampened by confounding effects from the remainder of the protein sequence.

**Fig 3.**
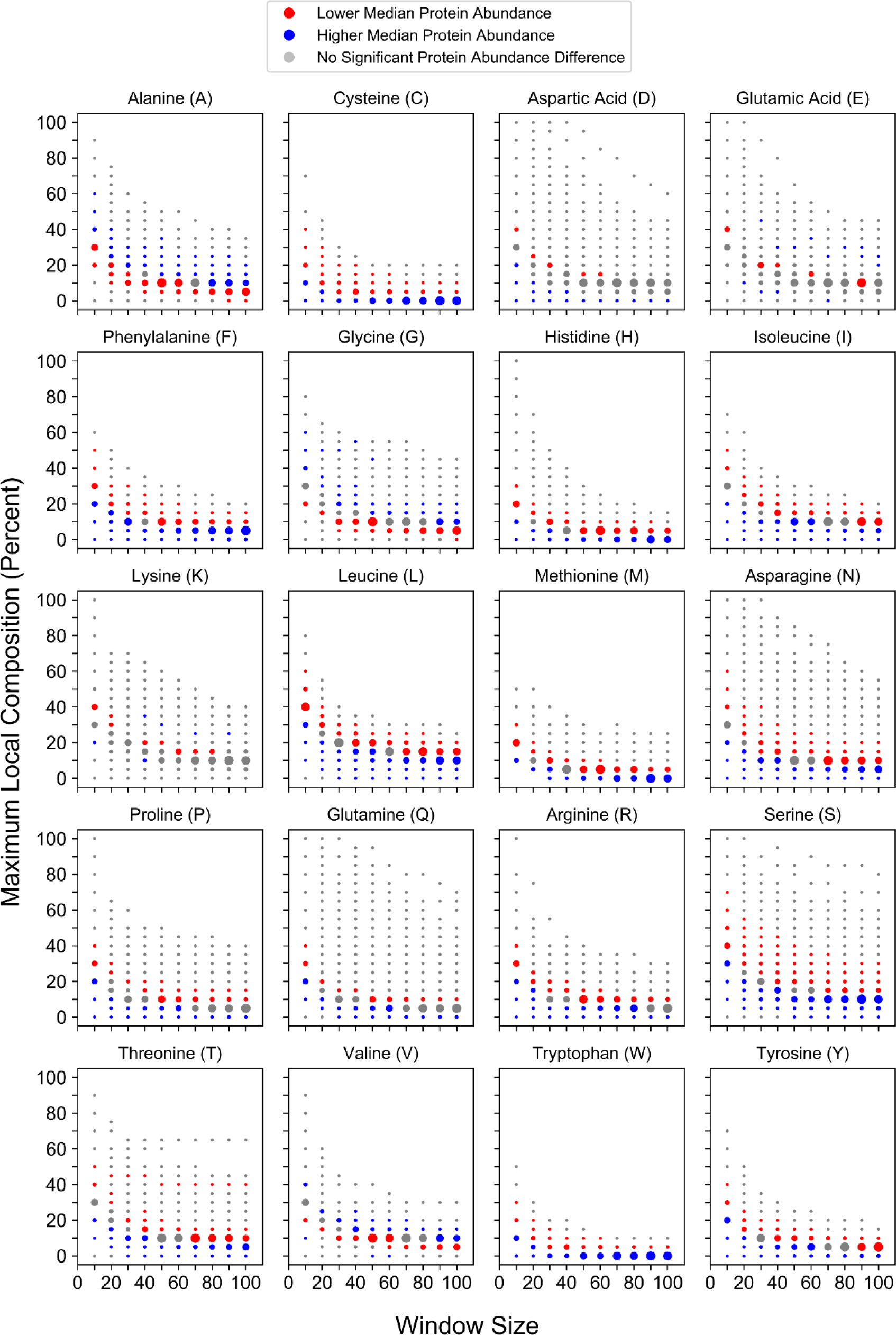
Maximum local amino acid composition affects protein abundance in a residue-specific manner. Local enrichment for individual amino acids correspond to changes in protein abundance. For each amino acid, protein abundance values corresponding to proteins partitioned into a given window size and percent composition bin were compared to values for all proteins of length ≥ the corresponding window size that were excluded from the bin. For this figure and subsequent, related figures, all colored points indicate statistically significant comparisons (Bonferroni-corrected *p* ≤ 0.05; see Methods). Bins corresponding to proteins with lower median values (relative to those of excluded proteins) are indicated in red, whereas proteins with higher median values (relative to those of excluded proteins) are indicated in blue. Comparisons lacking statistical significance are indicated in grey. Individual points are scaled within each subplot to reflect the sample sizes of proteins contained within each window size and percent composition bin.

Similar trends are observed when compositional enrichment is compared to protein half-lives (Fig 4). Compositional enrichment for the majority of amino acids (C, H, K, M, N, P, S, or T) is associated with lower protein half-life, whereas enrichment for A, G, I, or V is associated with higher protein half-life. Enrichment for F leads to an initial transition from lower to higher half-lives, while further enrichment leads to a transition back to lower half-lives. It is worth noting that similar trends were observed in an independent protein half-life dataset when the proteins were analyzed based on whole-protein amino acid composition [47], suggesting that maximum local composition is sufficient to detect associations between amino acid composition and half-life. Although for many amino acids the trends are readily apparent, the strength of the association between compositional enrichment and protein half-life appears to be slightly weaker than the association between compositional enrichment and protein abundance. This is likely due, at least in part, to limited proteome coverage (relative to the protein abundance dataset). However, a recent study also suggested that protein half-life is strongly affected by factors other than sequence characteristics [48], which would likely further dampen trends observed between compositional enrichment and protein half-life. Finally, protein half-life is generally less-conserved than protein abundance [49], perhaps suggesting that specific relationships between conserved sequence features and protein half-life may not be particularly strong. Therefore, it is rather surprising that we observe the indicated trends in spite of these limitations, and could suggest that half-life is more strongly influenced by local composition than particular primary sequence motifs.

**Fig 4.**
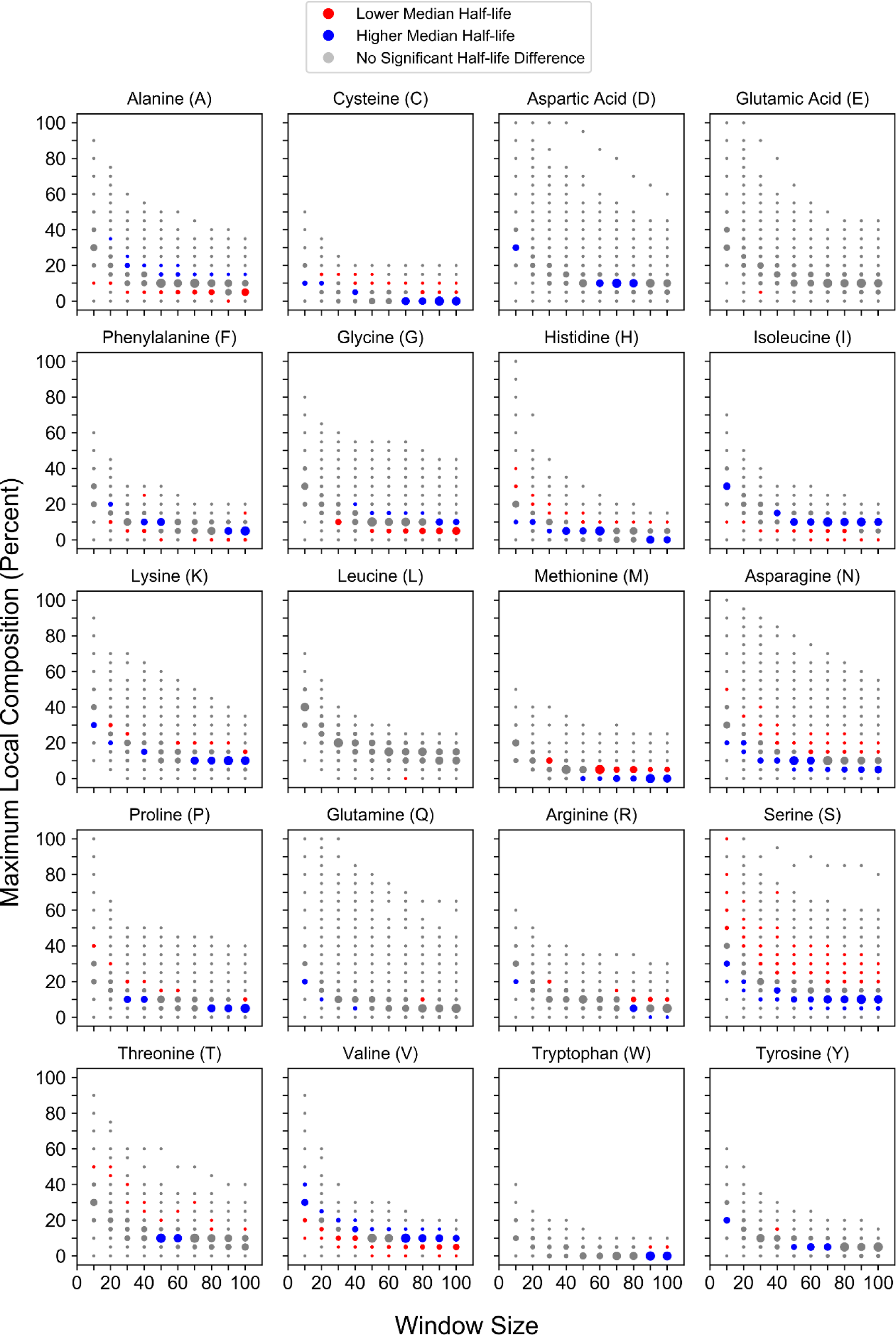
Maximum local amino acid composition affects protein half-life in a residue-specific manner. Local enrichment for individual amino acids correspond to composition-dependent changes in protein half-life.

Direct measurement of protein synthesis rates is more experimentally challenging. Consequently, proteome-wide coverage for experimentally-derived translation efficiency remains substantially lower than coverage for protein abundance and half-life. The normalized translation efficiency (nTE), a reported metric of translation elongation efficiency [50], is based on codon usage frequencies and tRNA gene copy numbers, allowing for calculation of translation efficiency for the entire proteome. Therefore, we first examined relationships between local compositional enrichment and calculated translation elongation efficiency. nTEs were calculated for whole-protein sequences using the corresponding coding region on mRNA transcripts (see Methods). Translation efficiency is strongly dependent on the locally-enriched amino acid (Fig 5). For the majority of amino acids (C, D, E, F, H, I, K, L, M, N, P, Q, R, or Y), local enrichment is associated with significantly lower median nTEs suggesting that, as a single class, proteins with local compositional enrichment tend to be translated relatively inefficiently. Proteins with domains enriched in S, T, or W are generally associated with significantly lower median nTEs, although proteins with very high S, T, or W enrichment are associated with significantly higher median nTEs. However, proteins with domains enriched in A, G, or V residues are consistently associated with significantly higher median nTEs, suggesting that these proteins may be translated relatively efficiently. Remarkably, nearly identical trends are observed between local compositional enrichment and the experimentally-derived protein synthesis rates reported for a limited proteome despite a substantial reduction in sample size (*n*= 1115) ([51]; Fig S1), suggesting that nTE can serve as a good surrogate for overall protein synthesis efficiency. Collectively, these results indicate that local amino acid enrichment is associated with differences in protein production rates in a composition-dependent manner.

**Fig 5.**
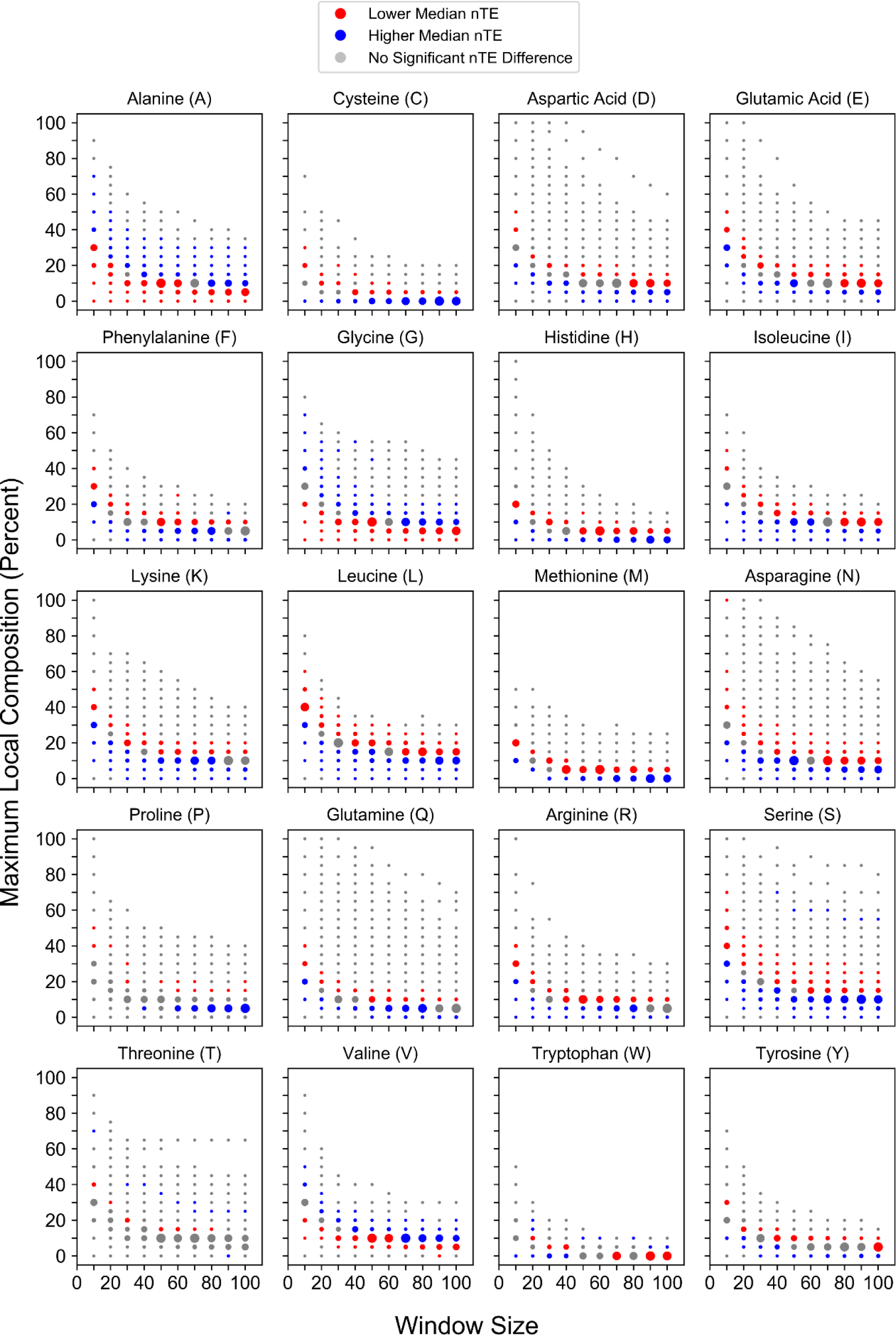
Maximum local amino acid composition affects nTE in a residue-specific manner. Local enrichment for individual amino acids correspond to composition-dependent changes in nTE.

For most amino acids, we noticed a remarkable correspondence in the trends for translation efficiency, protein abundance, and protein half-life, despite the fact that these values are derived from entirely different methods and experiments. For example, local enrichment for many amino acid types is associated with low nTE values, low protein abundance, and low protein half-life (Table 1). While translation efficiency and protein degradation rate are largely functionally independent in cells, protein abundance depends, at least in part, on both translation efficiency and protein half-life [44]. This may suggest that protein abundance for these proteins is limited in cells by a combination of poor translation efficiency and rapid degradation rate. In contrast, local enrichment for some amino acids is associated with high protein abundance also tended to have higher nTE values and higher half-lives, perhaps suggesting that high protein abundance for these proteins is achieved by a combination of efficient translation and poor degradation.

**Table 1:** The life-cycle of proteins with high local composition of individual amino acids involves the coordinated regulation of translation efficiency, protein abundance, and protein half-life. For each amino acid, trends in median values for nTE, protein abundance, and half-life upon enrichment (i.e. approaching higher percent compositions) of the given amino acid are indicated. “Higher” indicates that proteins in larger percent composition bins tend to have a larger median value compared to all other proteins, while “Lower” indicates that proteins in larger percent composition bins tend to have a larger median value compared to all other proteins. “Mixed” indicates amino acids for which multiple transitions are observed upon progressive compositional enrichment. Datasets without clear, statistically significant transition thresholds are also indicated (“n.s.”). Colors correspond to the colors used in Figs 3-5.

| <u>Amino Acid</u> | <u>nTE</u> | <u>Abundance</u> | <u>Half-life</u> |
| --- | --- | --- | --- |
| A | Higher | Higher | Higher |
| C | Lower | Lower | Lower |
| D | Lower | Lower | n.s. |
| E | Lower | Mixed | n.s. |
| F | Lower | Lower | Mixed |
| G | Higher | Higher | Higher |
| H | Lower | Lower | Lower |
| I | Lower | Lower | Higher |
| K | Lower | Mixed | Lower |
| L | Lower | Lower | n.s. |
| M | Lower | Lower | Lower |
| N | Lower | Lower | Lower |
| P | Lower | Lower | Lower |
| Q | Lower | Lower | n.s. |
| R | Lower | Lower | n.s. |
| S | Mixed | Lower | Lower |
| T | Mixed | Lower | Lower |
| V | Higher | Higher | Higher |
| W | Mixed | Lower | n.s. |
| Y | Lower | Lower | n.s. |

### Compositional Enrichment Exerts Biologically-Relevant Effects in the Absence of Classical Low-Complexity, Statistically-Biased, and Homopolymeric Domains

An important advantage of approaching LCDs from a composition-centric perspective is the ability to examine relationships between amino acid composition and protein outcomes without appealing to pre-defined thresholds of statistical amino acid bias [11] or sequence complexity [1,10], which may not reflect biologically-relevant thresholds. Indeed, the transitions observed in the median translation efficiencies, protein abundances, and protein half-lives often occur at surprisingly mild levels of compositional enrichment, suggesting that these effects may be observed even in the absence of classically-defined statistically-biased or low-complexity domains.

Statistical amino acid bias conceptually parallels our investigation of compositional enrichment, and has been used to investigate the functions of proteins with statistically-biased domains [11,12]. To examine whether compositional enrichment may exert biologically-relevant effects on protein metabolism independently of statistically-biased domains, a conservative bias threshold was employed to define statistically-biased domains using previously developed methodology [12] (also, see Methods). Proteins with statistically-biased domains were then filtered from the yeast proteome (*n* = 866 statistically-biased proteins for the yeast translated proteome of sequences ≥ 30 residues in length). However, even in the absence of statistically-biased domains, compositional enrichment resulted in robust trends in translational efficiency, protein abundance, and protein half-life that re-capitulated those originally observed (Figs S2-S4). This suggests that compositional enrichment affects protein metabolism at thresholds preceding those required for classification as statistically-biased by alternative methods.

The SEG algorithm, by default, employs substantially more relaxed criteria when classifying protein domains as low-complexity [1]. Indeed, of the 5,901 proteins of length ≥30 amino acids in the translated ORF proteome, 4,147 proteins contain at least one LCD, which is consistent with previous estimates [3]. Nevertheless, despite a large reduction in proteome size, many of the trends in protein metabolism are discernible even when all proteins with a SEG-positive sequence are filtered from the proteome (Figs S5-S7). This suggests that compositional enrichment exerts biologically relevant effects even among non-LCD-containing proteins.

Proteins containing homopolymeric amino acid repeats (defined as five or more identical amino acids in succession), were recently reported to have lower translation efficiency, lower protein abundance, and lower protein half-life when compared to proteins without homopolymeric repeats [36]. Proteins with homopolymeric repeats are expected to be disproportionately common among compositionally enriched domains, raising the possibility that the trends observed in the present study have been mis-attributed to compositional enrichment alone. To examine this possibility directly, the relationship between compositional enrichment and nTE, abundance, and half-life was re-evaluated for a filtered proteome that excludes all proteins containing at least one homopolymeric repeat (*n* = 755 proteins excluded). While exclusion of these proteins preferentially reduces the sample sizes at higher compositional enrichment percentages, the absence of homopolymeric repeat proteins has little effect on the trends in nTE, abundance, and half-life as a function of compositional enrichment (Figs S8-S10). This does not definitively rule out the possibility that homopolymeric repeats may, in some way, specifically affect translation efficiency, abundance, and half-life. However, since homopolymeric repeats *per se* are not absolutely required, the effects of homopolymeric repeats may instead be explained simply by local compositional enrichment.

Homopolymeric repeats are effectively short sequences of maximum possible single-amino acid density. However, the length of homopolymeric repeats is often, by necessity, arbitrarily defined. It is possible that very short sequences approaching homogeneity (i.e. near-homopolymeric sequences) could have similar effects. Indeed, the compositions corresponding to transition points in nTE, abundance, and half-life display some degree of sequence dependence – there is an inverse relationship between maximum local compositions required for transitions in nTE, abundance, and half-life and the window size used to scan protein sequences. Therefore, to ensure that the effects observed across window sizes are not due to very short sequences of high amino acid density, all sequences with a 10-amino acid window with at least 50% of any single amino acid were removed from the proteome. Notably, this includes proteins containing a canonical homopolymeric repeat, as well as an additional 3,479 proteins with short segments of high single amino acid density. Despite these considerably relaxed criteria and the dramatic reduction in whole-proteome sample size, many of the trends are still discernible (Figs S11-S13), particularly for the datasets with larger starting sample sizes (nTE and protein abundance). This indicates that the trends observed at larger window sizes are not solely due to the effects of very short sequences of high single amino acid density embedded within a larger window.

Collectively, these results suggest that compositional enrichment affects translation efficiency, protein abundance, and protein half-life at thresholds preceding those required for classification as low-complexity or statistically-biased by traditional methods. It is worth noting that in the course of eliminating proteins with classically-defined low-complexity, statistically-biased, or homopolymeric domains, proteins with multiple distinct domains strongly enriched in different amino acid types, or with single domains strongly enriched in more than one amino acid, are eliminated from the proteome before re-evaluation. Therefore, the trends in protein metabolism observed upon enrichment of a given amino acid are not due to confounding effects of domains strongly enriched in other amino acids occurring within the same protein sequences.

### Local Compositional Enrichment Influences Protein-Protein Interaction Promiscuity in a Residue-Specific Manner

Local enrichment of a single amino acid can dramatically influence the physicochemical properties of a given protein domain [23]. In a cellular context, these physicochemical properties likely influence interactions between proteins and surrounding molecules, including other proteins.

To examine whether local compositional enrichment affects protein-protein interactions, we explored relationships between enrichment for each of the amino acids and protein-protein interaction promiscuity (defined as the number of unique interacting partners per protein).Proteins found in a range of high-percent composition-bins for most amino acids (A, D, E, G, K, N, P, Q, R, and V) are associated with significantly more interacting partners relative to all other proteins (Fig 6), suggesting that these domains are relatively promiscuous. Additionally, proteins with mild enrichment for select hydrophobic residues (I, L, and M) are generally associated with more interacting partners, although fewer comparisons reach statistical significance (grey dots). These results are consistent with previous reports that, as a single class, proteins with LCDs or homopolymeric repeats tend to have more protein-protein interaction partners [16,36]. However, proteins in a range of high-percent composition-bins for each of the aromatic residues (F, W, and Y) are associated with significantly fewer interacting partners relative to other proteins, suggesting that aromatic residues tend to lack the interaction promiscuity observed at higher percent compositions for other amino acids. Furthermore, proteins with moderate to high local C content and proteins with extremely high maximum local S or T content are also associated with significantly fewer interacting partners relative to other proteins, suggesting that these domains are relatively non-promiscuous as well. This is particularly interesting, given that these effects were not observed upon enrichment for other polar residues. Again, this highlights the potential pitfall of grouping amino acids with related physicochemical properties into a single category.Collectively, these results indicate that protein-protein interaction promiscuity varies for proteins with high compositional enrichment in a residue-specific manner.

**Fig 6.**
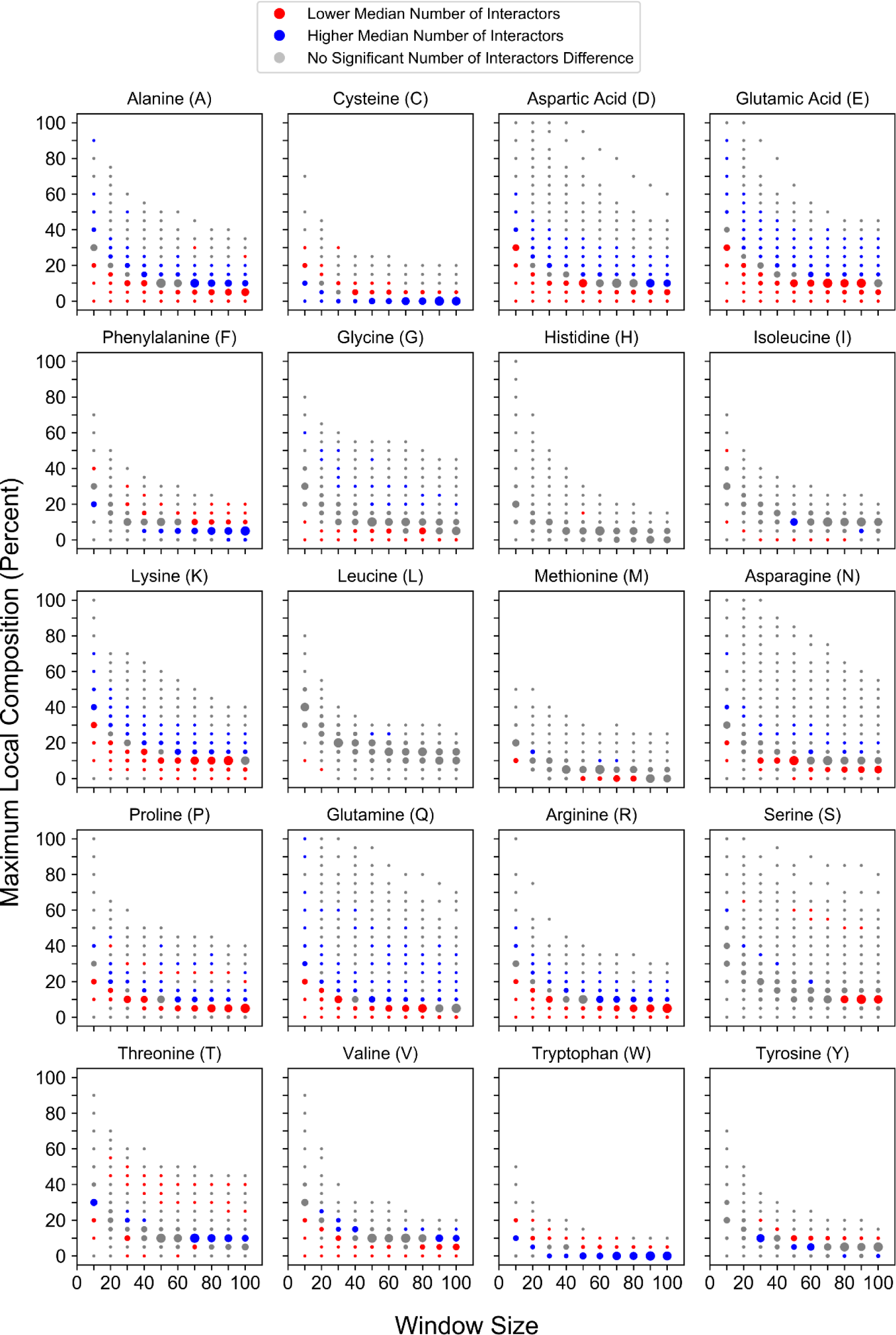
Maximum local amino acid composition affects protein-protein interaction promiscuity. Local enrichment for individual amino acids correspond to composition-dependent changes in the number of unique protein-protein interaction partners.

### Proteins with High Compositional Enrichment Can Fulfill Overlapping or Specialized Molecular Roles in the Cell

Previous studies have attempted to associate proteins containing LCDs, statistically-biased domains, and homopolymeric repeats with particular cellular functions [12,16–18,33,36]. However, one important consideration when inferring relationships between proteins with LCDs and cellular functions, for example, is the prevalence of proteins with multiple LCDs [3], and of LCDs strongly enriched in more than one amino acid type [11,14,18,35]. Therefore, attempts to associate cellular functions to specific LCD types, without controlling for other LCDs within the same protein sequences, risk mis-attributing functions to unrelated protein features [12,14,33,35]. While multiple LCDs within the same protein (or multiple amino acid types enriched within the same LCD) may cooperate to generate novel structures or functions, this complicates interpretation of the role of each individual amino acid type within LCDs.Furthermore, because some types of LCDs are more common than others, general attempts to associate cellular functions with LCDs, statistically-biased domains, or homopolymeric repeats likely reflect the functions associated with only the most common types when considered as a single, unified class [16,36]. Therefore, definitive assignment of cellular functions to each individual class of LCD necessitates exclusion of proteins with other types of LCDs.

In order to minimize possible confounding effects introduced by proteins with multiple regions enriched in different amino acid types, a modified version of the initial calculation performed by the SEG algorithm (namely, the Shannon entropy; see Methods) was employed to define proteins with only a single type of compositionally-enriched domain (CED). In an effort to incorporate our results (which indicate that compositional enrichment can exert biologically-relevant effects at compositions preceding the SEG algorithm threshold) into our definition of single-CED proteins, percent composition bins for which at least 75% of the residing proteins contained a SEG-positive sequence (as defined above) were pooled to generate a single list of CED-containing proteins for each amino acid. Proteins that contain multiple types of CEDs were then removed from the dataset, resulting in a non-redundant set of proteins with only one type of CED. Importantly, this method captures the exclusion of proteins containing more than one type of CED, as well as proteins with CEDs strongly enriched in more than one amino acid type.

Gene Ontology (GO) term analysis was performed separately for each window size within each single-CED category. For each type of CED, there is strong overlap in the enriched GO terms across the range of window sizes, suggesting that the associations between functions and residue-specific CEDs are not strongly length-dependent at this scale. Therefore, for simplicity of interpretation, significantly enriched GO terms for each window size were pooled to generate a single non-redundant list of enriched GO terms for each CED type.

Removal of proteins with multiple types of CEDs reveals a remarkable degree of specialization for CEDs of different types (Fig 7, and Table S2, which is often not observed for CEDs when considered as a single category or when multi-CED proteins are not excluded. For example, L-rich proteins are predominantly associated with functions at the ER and vacuole membranes, whereas I-rich proteins are more strongly associated with carbohydrate transport at the plasma membrane. A-rich proteins are associated with a variety of processes or cellular components, including translation, protein kinase activity, the cell wall, and carbohydrate/alcohol catabolism. N-rich proteins are strongly associated with functions related to transcription, whereas Q-rich proteins are associated with endocytosis and other cytoplasmic processes.Finally, although yeast cell wall proteins are often radically S/T-rich, after controlling for co-enrichment of S and T in the same proteins, S-rich proteins are more strongly associated with membrane-related processes (cell wall, cellular bud tip, cellular bud neck, mating tip projection, etc.), protein kinase activity, and transcription, whereas T-rich proteins tend to be associated with nucleic acid binding and helicase activity, with fewer associations with membrane-related processes. Therefore, specialized functions emerge even among commonly grouped amino acids, after controlling for the presence of multiple CEDs within the same proteins.

**Fig 7.**
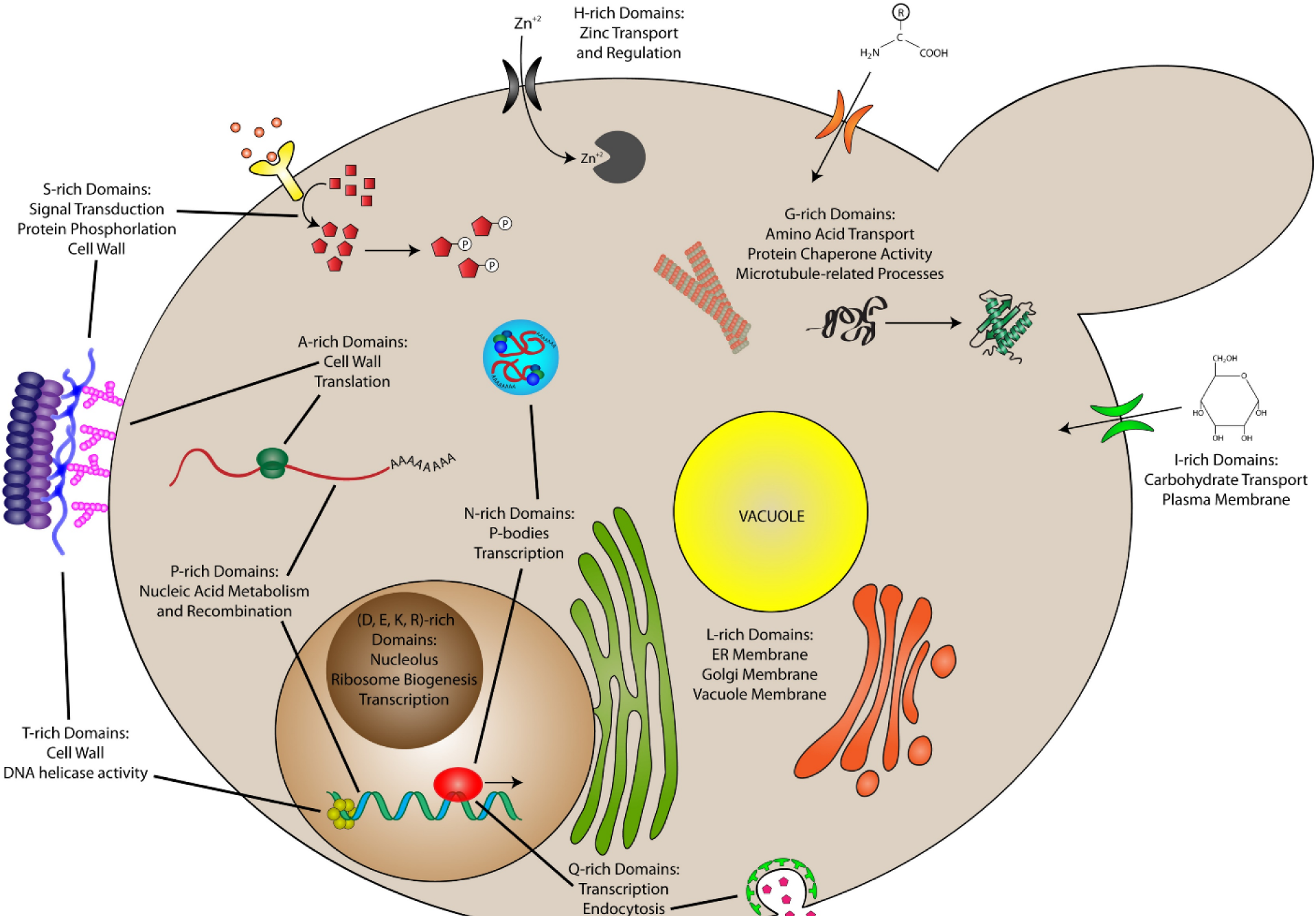
Cell model depicting overlapping and distinct functions of CEDs.

Furthermore, CEDs enriched in some amino acids share functions despite the removal of multi-CED proteins, suggesting some degree of co-specialization. For example, D-, E-, and K-rich CEDs were each associated with functions in the nucleus/nucleolus, including ribosomal RNA processing, nucleic acid binding, and transcription. Intriguingly, intrinsically disordered domains with opposite net charges (along with other charged macromolecules such as nucleic acids and polyADP-ribose) can drive phase separation or complex coacervation in the nucleus [52–54]. It is possible that these domains, along with nucleic acids and other polyionic molecules, may participate in nuclear processes via dynamic electrostatic association with these or other membraneless assemblies. By contrast, H-rich CEDs are associated with processes related to zinc ion transport and regulation. There were no GO terms significantly associated with R-rich CEDs. However, compositional enrichment for R appears to be constrained, as evidenced by the sharp decline in the number of proteins with R-rich domains toward higher maximum local percent compositions (see Fig 2), which may be further impacted by the removal of proteins with other types of CEDs.

In summary, when examined as separate classes, different types of CEDs can have overlapping or specialized roles in the cell.

### Compositional Enrichment Corresponds to Preferential Localization to Specific Subcellular Compartments

The molecular specialization observed for CEDs indicates that proteins with enrichment of particular residues may localize to particular subcellular compartments in order to execute their specialized functions. Furthermore, protein quality control factors can differ between subcellular compartments (for review, see [55]), which may contribute to composition-dependent differences in protein metabolism (see Fig 2). Therefore, we applied a bottom-up approach to infer the composition profiles associated with the major subcellular compartments (see Methods).

Largely aqueous subcellular compartments are almost exclusively associated with proteins containing domains enriched in charged residues, polar residues, and proline (Fig 8; see also Figs S13-S21). However, differences in compositional enrichment profiles are apparent even among related aqueous compartments. For example, significant associations with charged, Q, or N residues reach more extreme percent compositions in the nucleus, whereas as significant associations with P enrichment reach higher percent compositions in the cytoplasm. By contrast, the highly membraneous internal organelles (i.e. the endoplasmic reticulum and Golgi apparatus) are predominantly associated with enrichment of hydrophobic or aromatic residues. The yeast vacuole is also associated with composition profiles resembling those of membraneous compartments, with additional weaker associations with S and C enrichment.Few weak associations are observed for mitochondria. The yeast cell wall is strongly associated with S enrichment (likely related to its ability to be glycosylated), with additional moderate associations with T and A enrichment, and a weak association with mild V enrichment. As expected, the plasma membrane is also associated with enrichment for a variety of hydrophobic and aromatic residues. However, the plasma membrane is also significantly associated with enrichment of a select subset of polar residues (namely C, G, S, and T), further corroborating the specialized roles observed for these CEDs at the outer membrane. Indeed, G-rich CEDs are significantly associated with amino acid transport (see Table S2, and S-or T-rich CEDs of the plasma membrane could have overlapping functions or interactions with S-and T-rich CEDs of the cell wall. Together, these observations indicate that subcellular compartments may tolerate or prefer proteins with specific types of CEDs.

**Fig 8.**
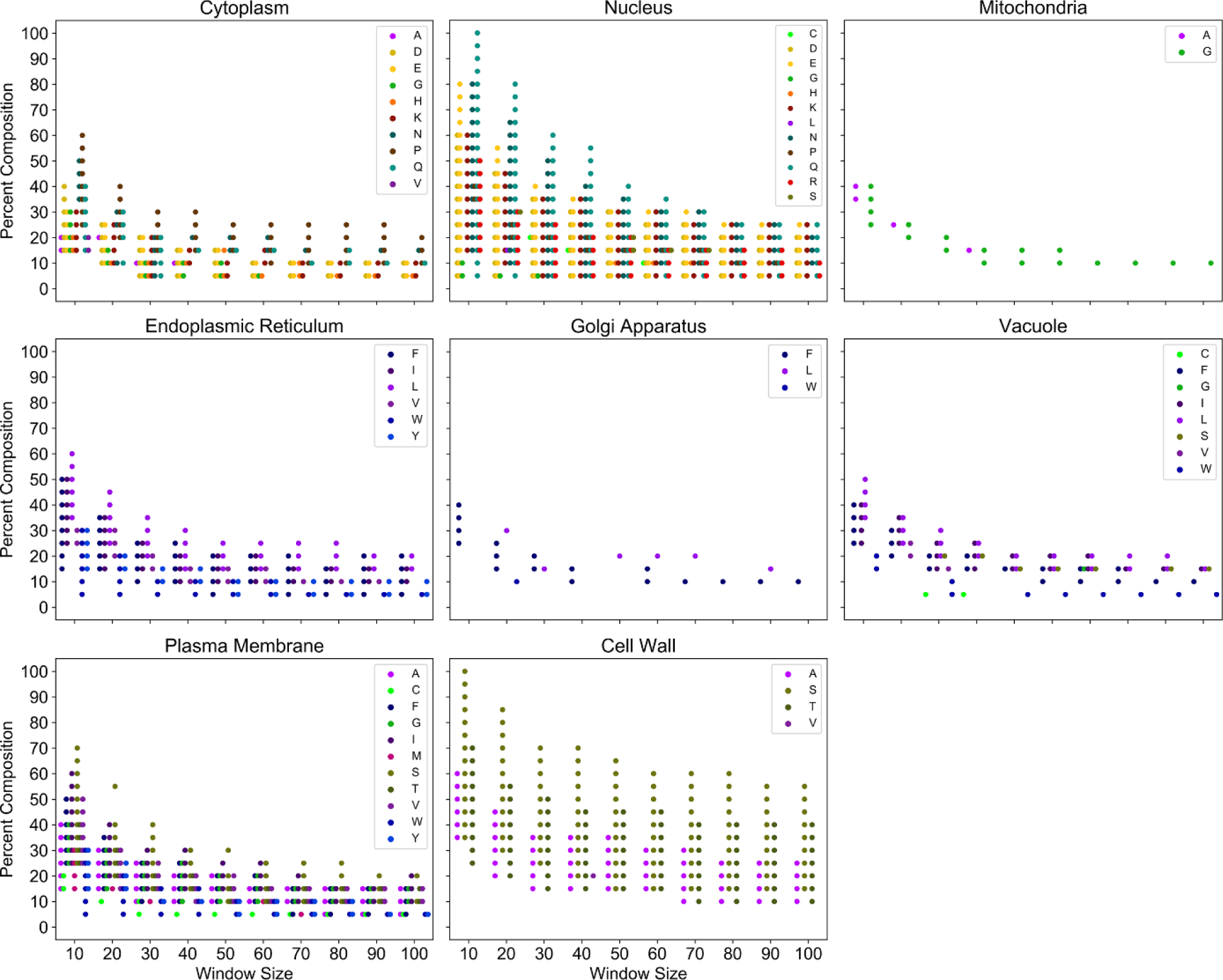
Compositional enrichment profiles associated with major subcellular compartments. All plotted points indicate protein sets for which association with the indicated subcellular compartment is statistically significant (Fisher’s exact test, with Bonferonni-corrected *p* < 0.05). Warm colors (reds, oranges, and yellows) correspond to charged residues. Green colors indicate polar residues. Cool colors (purples and blues) correspond to hydrophobic and aromatic residues respectively.

### Components of Stress Granules and Processing Bodies Possess Shared and Unique Compositional Features

Recent observations indicate that a variety of Q/N-rich and G-rich domains can form highly dynamic protein-rich droplets in aqueous environments [24–29], a process referred to as liquid-liquid phase separation. These types of LCDs are prevalent among components of membraneless organelles such as stress granules and P-bodies [22]. Furthermore, stress granules and P-bodies share many properties with protein-rich liquid droplets formed *in vitro*, suggesting that the fundamental biophysical properties of these domains are related to the formation of membraneless organelles *in vivo*. However, while amino acid composition is acknowledged as a critical determinant of this behavior, the precise compositional requirements associated with membraneless organelles remain largely undefined.

Therefore, we also applied our bottom-up approach to infer the compositional enrichment profiles associated with protein components of stress granules and P-bodies (as defined in [56]). Stress granules and P-bodies have overlapping protein constituents and can exchange protein components [57,58], suggesting that they are closely related yet distinct organelles. Accordingly, we observe both shared and unique features in the composition profiles associated with stress granule and P-body proteins (Fig 9). As expected, both stress granules and P-bodies are strongly associated with proteins containing Q-rich or N-rich domains. For example, minimum Q or N compositions significantly associated with stress granules range from ∼15-100% at small window sizes (≤30 amino acids) and ∼10-30% at large window sizes (≥80amino acids), although these values vary slightly depending on window size and residue. Similarly, minimum Q or N compositions significantly associated with P-bodies range from ∼15-100% at small window sizes and ∼10-40% at larger window sizes.

**Fig 9.**
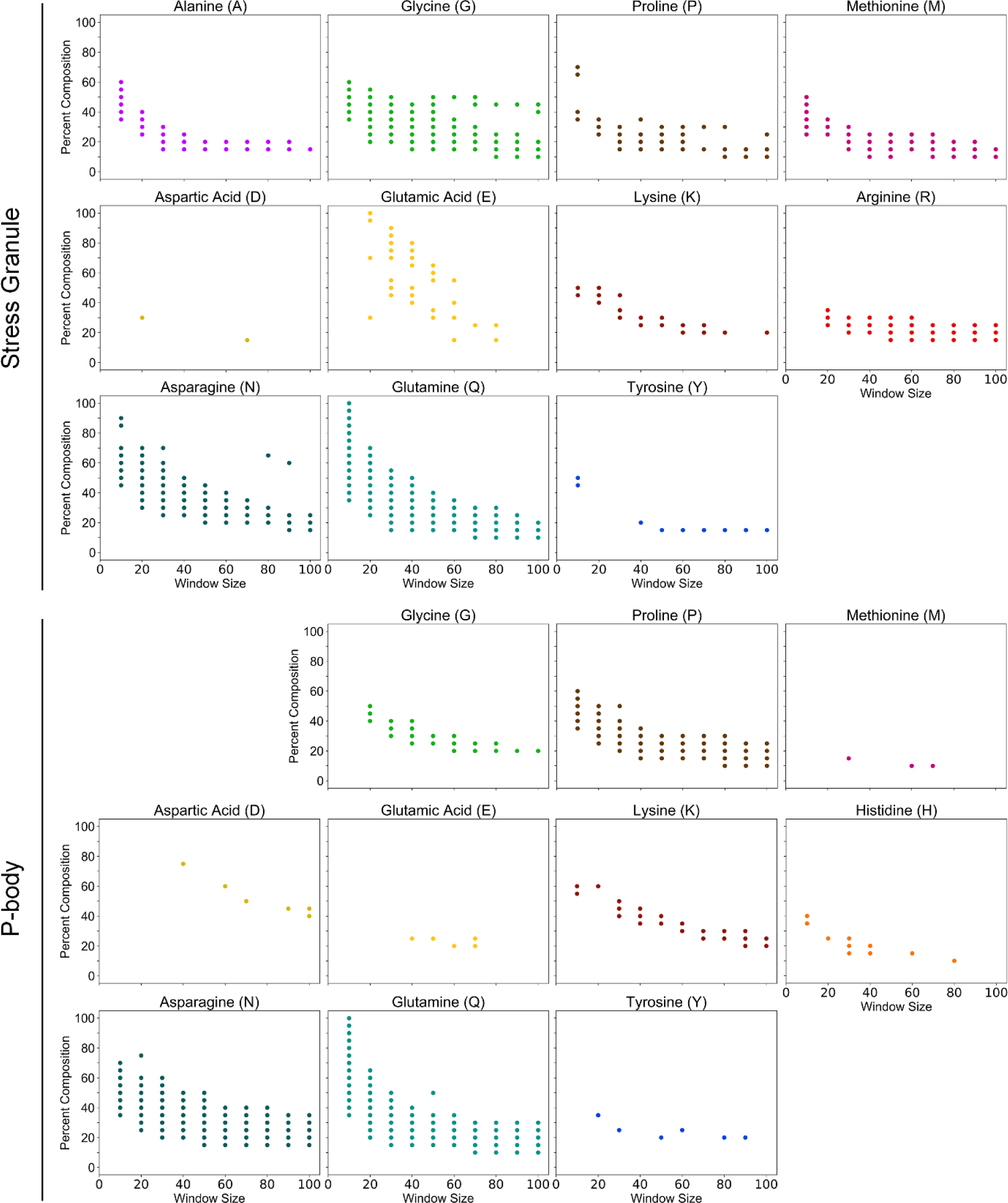
Composition profiles associated with membraneless organelles. All colored points indicate minimum percent composition thresholds for which components of stress granules (A) or P-bodies (B) are significantly enriched (*p* < 0.05). Only amino acids for which significant enrichment of stress granule or P-body proteins was observed in at least two composition bins are shown. For greater sensitivity, plots were generated using uncorrected *p*-values. Therefore, any individual point should be viewed with some skepticism: however, the presence of multiple consecutive significant points within each window size suggest that the observed trend is likely not an artifact of multiple hypothesis testing (see Fig S22 for analogous plots with Bonferroni-corrected *p*-values).

In addition to the commonly appreciated link between stress granule/P-body components and Q/N-rich domains, we identify and define a variety of previously unappreciated compositional features common to stress granule and P-body components. Components of both stress granules and P-bodies are strongly associated with P-rich domains, weakly associated with K-rich domains, and very weakly (yet significantly) associated with Y-rich domains.Furthermore, while both stress granules and P-bodies are associated with proteins containing G-rich domains, stress granule components are associated with a much broader range of G enrichment, suggesting that G enrichment may be a more characteristic feature of stress granules than P-bodies.

Additionally, some compositional features are unique to either stress granules or P-bodies. For example, stress granule constituents are significantly associated with A-rich, M-rich, E-rich, and R-rich domains, whereas P-body constituents exhibit little or no preference for these compositional features. By contrast, P-body components are weakly associated with H-rich domains, whereas stress granule components are not enriched among proteins containing H-rich domains.

To our knowledge, this represents the first attempt to systematically define the range of amino acid compositions associated with membraneless organelles such as stress granules and P-bodies. These observations suggest that components of related, membraneless organelles have overlapping yet distinct compositional preferences. It is possible that shared compositional features facilitate the physical interactions between stress granules and P-bodies and allow for the exchange of components, while differences in compositional features facilitate their ability to function as independent organelles.

## Discussion

Protein domains categorized as low-complexity, statistically-biased, or homopolymeric encompass broad, heterogeneous classes of sequences with diverse physical properties and cellular functions. These domains can play important roles in normal and pathological processes. However, challenges in categorizing proteins on the basis of sequence complexity or statistical bias have thus far precluded a complete, proteome-wide view of the effects of these domains on protein regulation and function. Here, we adopt an alternative, unbiased approach to examine proteome-wide relationships between local amino acid enrichment and the birth, abundance, functions, subcellular localization, and death of proteins. For nearly all amino acids, progressive local enrichment corresponds to clear transition thresholds with regard to translation efficiency, protein abundance, and protein half-life. Transition thresholds ubiquitously occurred at compositions preceding those required for classification as low-complexity or statistically-biased by traditional methods, indicating that our observed transition thresholds more closely reflect biologically-relevant composition criteria.

Protein sequences can range from perfectly diverse (i.e. a completely homogeneous mixture of amino acids with maximal spacing between identical amino acids) to lacking any diversity (i.e. homopolymeric sequences). While homopolymeric regions represent an extreme on this spectrum and can influence protein metabolism [36], classically defined homopolymeric regions are not absolutely required for these effects (see Figs S8-S10). Furthermore, many of the trends were discernible even upon removal of all proteins with a short stretch of high single-amino acid density. This suggests that compositional enrichment can affect protein metabolism even upon some degree of primary sequence dispersion (i.e. greater linear spacing between identical amino acids). Defining the limits of this dispersion may shed additional light on the relationship between amino acid composition and protein metabolism.

An advantage of assessing compositional enrichment (as opposed to sequence complexity) is the ability to distinguish the effects of compositional enrichment for each amino acid type. The nature of the trends in translation efficiency, protein abundance, and protein half-life depend on the amino acid enriched in the protein sequences, indicating that local enrichment of different amino acids can have opposite effects. This highlights a key limitation when considering low-complexity, statistically-biased, or homopolymeric domains as a single class – grouping domains composed of radically different amino acids effectively skews any trends observed toward those of the most common type and, in some cases, can completely mask the effects of less common low-complexity, statistically-biased, or homopolymeric domains. Furthermore, even grouping these domains on the basis of common physicochemical properties can introduce the same complication. This is exemplified by the non-aromatic hydrophobic amino acids; while I-rich, L-rich, and M-rich domains are associated with poor translation efficiency, low abundance, and rapid degradation rate, A-rich and V-rich domains are associated with high translation efficiency, high abundance, and slow degradation rate. Additionally, the cellular functions associated with domains enriched in hydrophobic residues tend to differ; L-rich domains are predominantly associated with the ER or vacuole membrane, whereas I-rich domains are predominantly associated with carbohydrate transport at the plasma membrane. Similarly, N-rich domains are strongly associated with transcription-related processes, whereas Q-rich domains are more strongly associated with endocytosis and other processes in the cytoplasm. While there is some overlap between these two groups, this suggests that domains enriched in remarkably similar amino acids may yet be favored for specialized roles in the cell.

Finally, a bottom-up application of our composition-centric algorithm to membraneless organelles provides the first step in defining the distinct compositional profiles associated with each type of organelle. We find that even closely related and physically interacting organelles are associated with discernible differences in compositional enrichment, which may relate to differences in their properties, regulation, and function *in vivo*.

While a great deal of attention is rightfully devoted to understanding relationships between primary amino acid sequence and protein fates (including folding, regulation, and functions), amino acid composition is increasingly believed to drive a variety of cellular and molecular processes. Here, we have developed an approach to examine the effects of local compositional enrichment for each of the canonical amino acids, in the absence of *a priori* assumptions or pre-defined thresholds. Our results provide a coherent, proteome-wide view of the relationships between compositional enrichment and the fundamental aspects of protein life cycle in a eukaryotic organism.

## Methods

### Composition-Based Proteome Scanning Algorithm

Yeast protein sequences were parsed using FASTA sequence parsing module from the Biopython package [59]. For each amino acid in the set of 20 canonical amino acids, each protein in the translated ORF proteome (latest release from the Saccharomyces Genome Database website, last modified 13-Jan-2015) was scanned using a sliding window of defined size (ranging from 10 to 100 amino acids, in increments of 10). The percent composition of the <u>a</u>mino <u>a</u>cid <u>o</u>f interest (AAoI) is calculated for each window, and the protein is sorted into bins based on the maximum percent composition achieved for the AAoI (ranging from 0 to 100 percent composition in 5 percent increments). Analyses were performed for all possible AAoI, window size, and percent composition combinations.

### Normalized Translation Efficiency (nTE)

Translation efficiency for each gene was estimated using the normalized translation efficiency (nTE) scale [50], which is based on tRNA gene copy number, codon-anticodon wobble base-pairing efficiency, and transcriptome-wide codon usage. However, the original nTE algorithm plots all nTE values for each codon to generate a separate translation efficiency profile for each gene. In order to condense translation efficiency information to a single value for each gene (in a manner analogous to the tRNA adaptation index; [60]), the geometric mean of nTE values across the transcript was calculated as

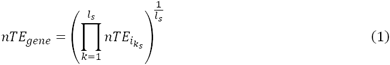

where *nTE*_*iks*_ represents the translation efficiency value of the *i*th codon defined by the *k*th triplet in nucleotide sequence *s*, and *l*_*s*_ represents the length of the nucleotide sequence excluding stop codons. Therefore, nTE values reported in the current study represent whole-gene nTE values. nTE analyses were performed using an in-house Python script.

### Shannon Entropy

The Shannon entropy of each sequence was calculated as

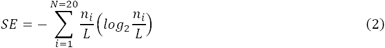

where *N* represents the size of the residue alphabet (*N* = 20, for the canonical amino acids), *n*_*i*_ represents the number of occurrences of the *i*th residue within the given sequence window of length *L*. For comparison with established measures of sequence complexity, we defined low-complexity domains by using the default window size (12 amino acids) and Shannon entropy threshold (SE ≤ 2.2bits) used in the first pass of SEG algorithm to initially identify LCDs [1,10].

In the SEG algorithm, the complexity state vector used to calculate the Shannon entropy is blind to the amino acid composition (i.e. the *n*_*i*_ values in Equation 2 are not attributed their respective amino acids). Therefore, when indicated, in order to distinguish LCDs on the basis of the predominant amino acid, sequences for which the SE ≤ 2.2bits and *n*_*AAoI*_ ≥ *n*_*max*_ within the complexity state (indicating that the AAoI is a major contributor to the sequence’s classification as an LCD) were assigned to the corresponding amino acid category (e.g. A-rich LCDs, C-rich LCDs, etc.). Single-LCD/CED proteins are proteins classified as LCDs or CEDs that do not appear on multiple amino acid-specific LCD/CED lists.

### Statistical Amino Acid Bias

Statistical amino acid bias was calculated as described in [12]. Briefly, the lowest probability subsequence for each protein was determined by exhaustively scanning proteins with window sizes ranging from 25 to 2500 amino acids. For each window, the subsequence bias probability (P_*bias*_) was defined as

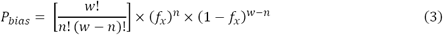

where *w* denotes the window size, *n* denotes the number of occurrences of the amino acid of interest within the subsequence, and *f*_*x*_ denotes the fraction of the amino acid of interest in the yeast translated proteome. The lowest probability subsequence for each protein is the subsequence with the lowest P_*bias*_.

A suitable threshold to define statistically-biased proteins within the yeast protein was determined as previously described [12], except that more relaxed criteria were used in order to include additional proteins with less extreme biases. Briefly, the P_*bias*_ corresponding to the lowest probability subsequence (P_*min*_) for each protein was plotted on a log-log plot against whole-protein sequence length. A line was fitted, then the y-intercept was decreased until only 15% of the proteome had P_*min*_ values below the line (previous analyses used a more stringent cutoff of 10% to define statistically-biased proteins). Additionally, a length-independent threshold was designated as the P_*min*_ value at which 15% of the proteome had smaller absolute P_*min*_ values. This threshold was used when it was less than the P_*min*_ threshold given by the length-dependent method to avoid unreasonably relaxed bias criteria for small protein sequences.Amino acid bias was calculated using values from the translated orf proteome only, and implemented via an in-house Python script with pre-computed look-up tables for computational efficiency.

### Homopolymeric and Near-Homopolymeric Sequences

Proteins containing homopolymeric sequences were defined simply as any protein with a subsequence of five or more contiguous residues of the same amino acid, as previously described [36]. Additionally, in order to explore more relaxed criteria, proteins with near-homopolymeric sequences were defined as any protein with a 10-amino acid subsequence containing at least 50% composition of any single amino acid.

### Protein Abundance and Protein Half-Life Data

Protein abundance values (in average number of molecules per cell per protein) were obtained from [43] (*n* = 5,391). Protein half-life data were obtained from [44]. For simplicity of interpretation, only proteins with unambiguous, non-zero half-life values (in minutes) were included in the datasets. Proteins listed on separate lines with identical half-life values were retained, whereas protein half-life measurements assigned to more than one protein on the same line were excluded (these were often highly homologous genes, suggesting that the half-life measurement could not be unambiguously assigned to one of the proteins). Furthermore, all proteins corresponding to “low-confidence” measurements were excluded (see [44] for criteria). *n* = 3,525 for the filtered half-life dataset.

### Statistics and Plotting

For all AAoI/window size/percent composition bins, the distribution of nTE, abundance, or half-life values for proteins included in the given bin was compared to the distribution of the respective values of all proteins excluded from the given bin. Statistical significance was estimated using a two-sided Mann-Whitney *U* test (also referred to as the Wilcoxon rank-sum test; refer to Supplemental Experimental Procedures from [42] for a detailed description and rationale). All reported *p*-values were adjusted within each window using the Bonferroni correction method for multiple hypothesis testing. All statistical tests were performed using modules available in the SciPy package with default settings, unless otherwise specified. All plots were generated using Matplotlib or Seaborn modules.

### Gene Ontology (GO) Term Enrichment Analyses

GO term enrichment tests were performed using the GOATOOLS package (version 0.7.9) [61] for each set of proteins contained in a given amino acid/window size/percent composition bin. For each test, the set of background proteins was defined as all proteins from the translated ORF proteome of sequence length greater than or equal to the given window size. All reported *p*-values were adjusted using the Bonferroni correction during GO term association. To evaluate the compositional enrichment profiles associated with GO terms related to subcellular compartments, we applied a minimum-threshold-scanning approach to all partitioned proteomes. For each AAoI, window size, and percent composition bin, all proteins with maximum local compositions greater than or equal to the current percent composition under consideration are pooled and evaluated for possible enriched GO terms. This effectively evaluates possible GO term enrichment iteratively with increasing maximum local composition criteria. GO term results were subsequently evaluated for significant enrichment of a single GO term describing each subcellular compartment (or two related GO terms, “outer membrane” and “plasma membrane”, in the case of the plasma membrane). *p*-values were further adjusted within each window size using the Bonferroni correction method.

Similar analyses were performed for the sets of experimentally-defined stress granule (*n*= 83) and P-body (*n* = 52) proteins [56]. Specifically, a minimum-threshold-scanning approach was applied to all partitioned proteomes. For each AAoI, window size, and percent composition bin, all proteins with maximum local compositions greater than or equal to the current percent composition under consideration are pooled. Significant enrichment of experimentally-defined stress granule or P-body proteins within each pool of proteins was evaluated using Fisher’s exact test (*p* < 0.05, adjusted within each window size using the Bonferroni correction method).

## Supporting Information

**Table S1. Maximum local composition values for each amino acid and window size combination for all translated yeast proteins.** For each protein in the translated ORF proteome, nTE, protein abundance, and protein half-life values are indicated. nTE was calculated according to the method described in [50]. Protein abundance and protein half-life values were reported in [43] and [49], respectively. Remaining columns contain the maximum local composition value (per 100) for each amino acid and window size combination for each gene.

**Table S2. Residue-specific CEDs are associated with unique cellular structures and processes.** All GO terms listed represent terms significantly associated with a set of residue-specific CEDs for at least one window size (Bonferroni-corrected *p* ≤ 0.05).

---

**Fig S1.**
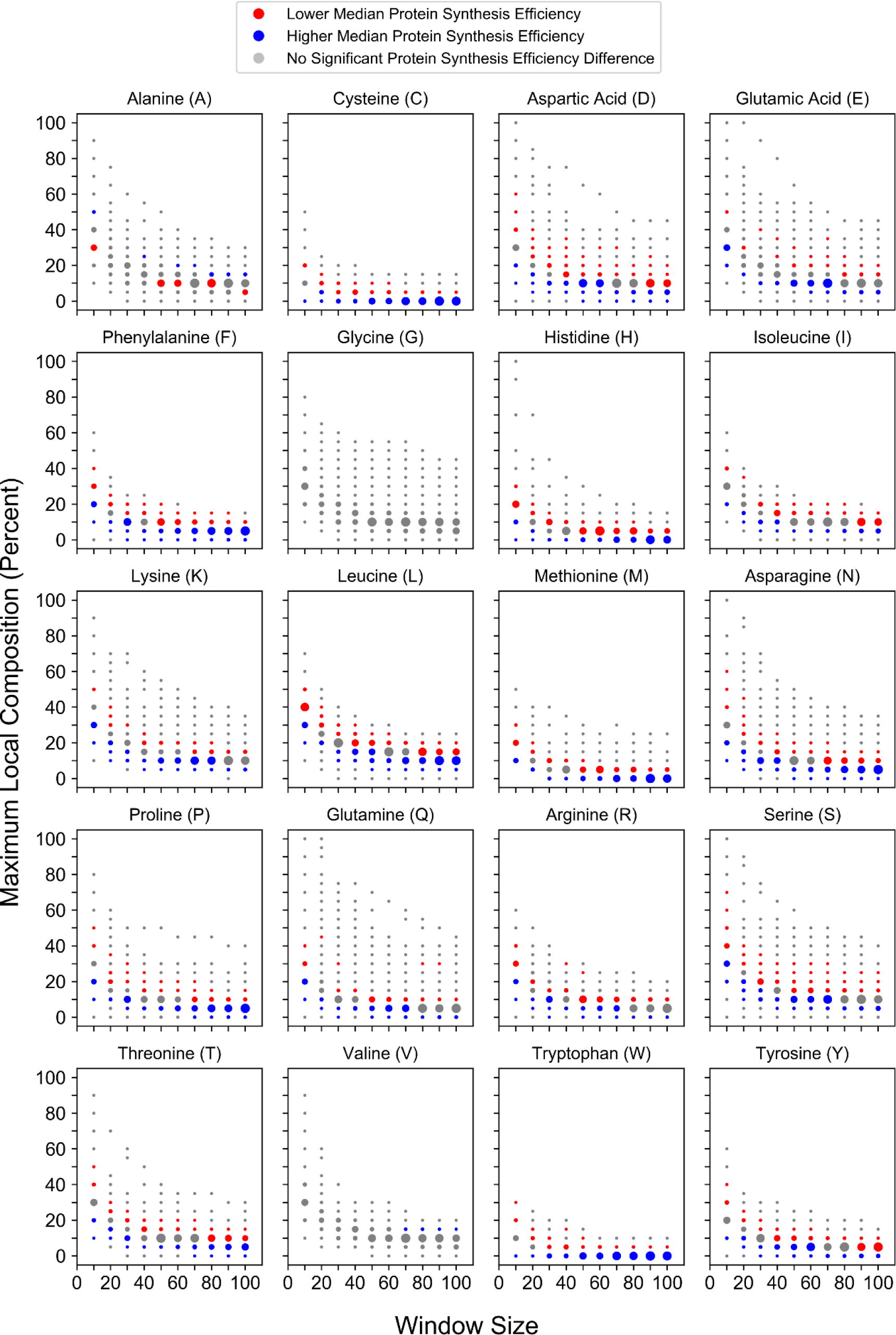
Maximum local amino acid composition affects protein synthesis efficiency in a residue-specific manner. Local enrichment for individual amino acids correspond to composition-dependent changes in experimentally-derived protein synthesis efficiency [51].

**Fig S2.**
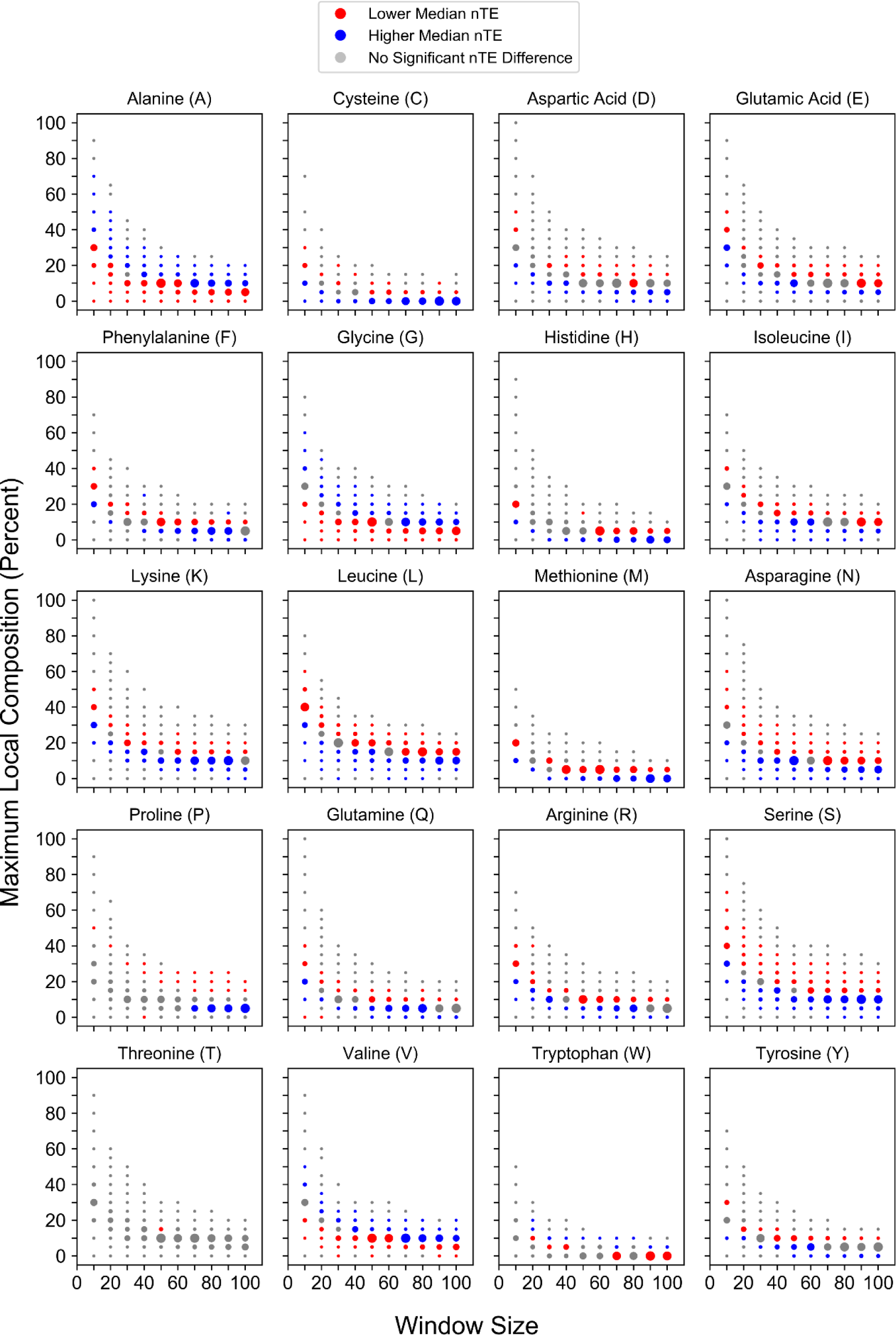
Local compositional enrichment affects the nTE of proteins lacking statistically-biased domains.

**Fig S3.**
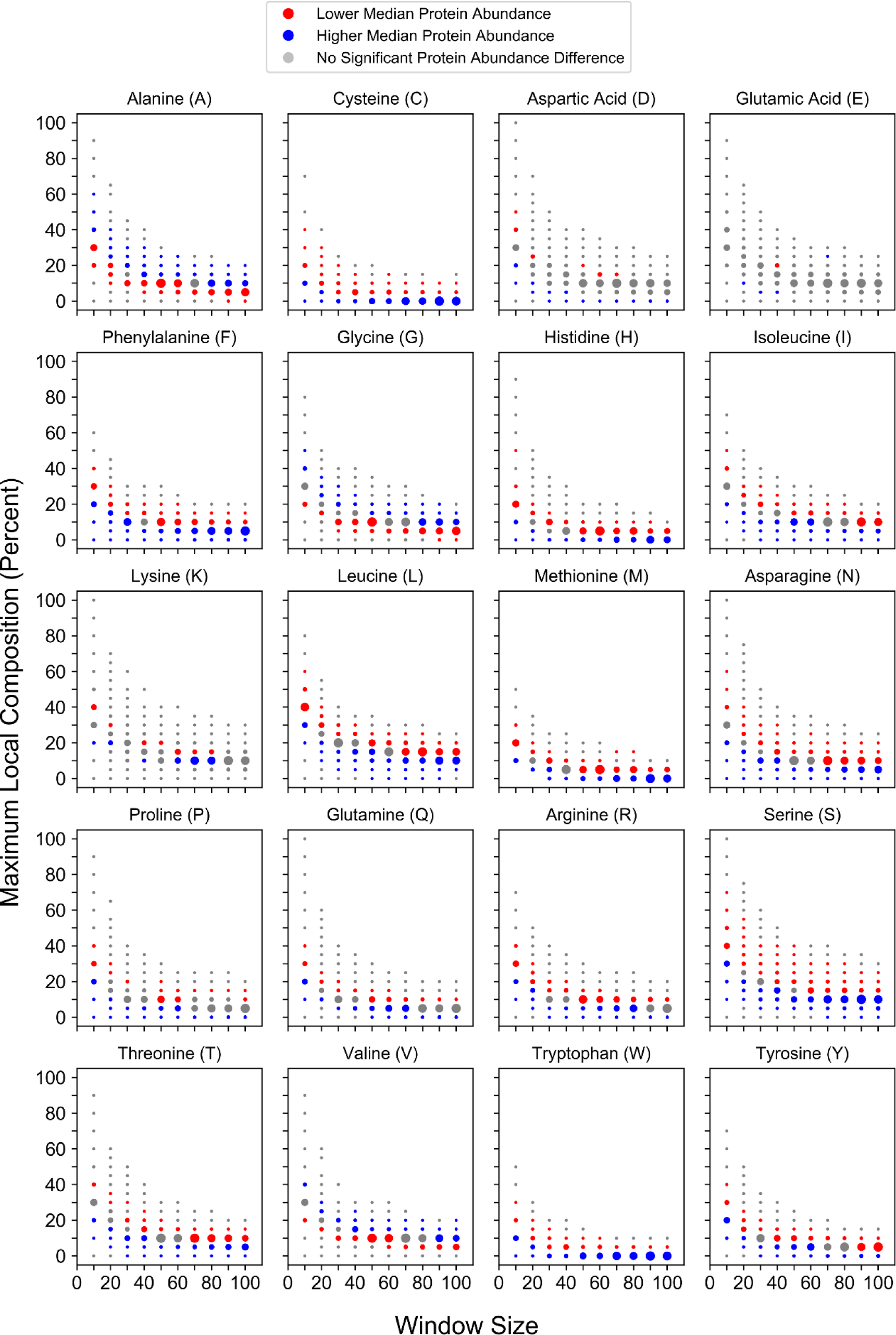
Local compositional enrichment affects the abundance of proteins lacking statistically-biased domains.

**Fig S4.**
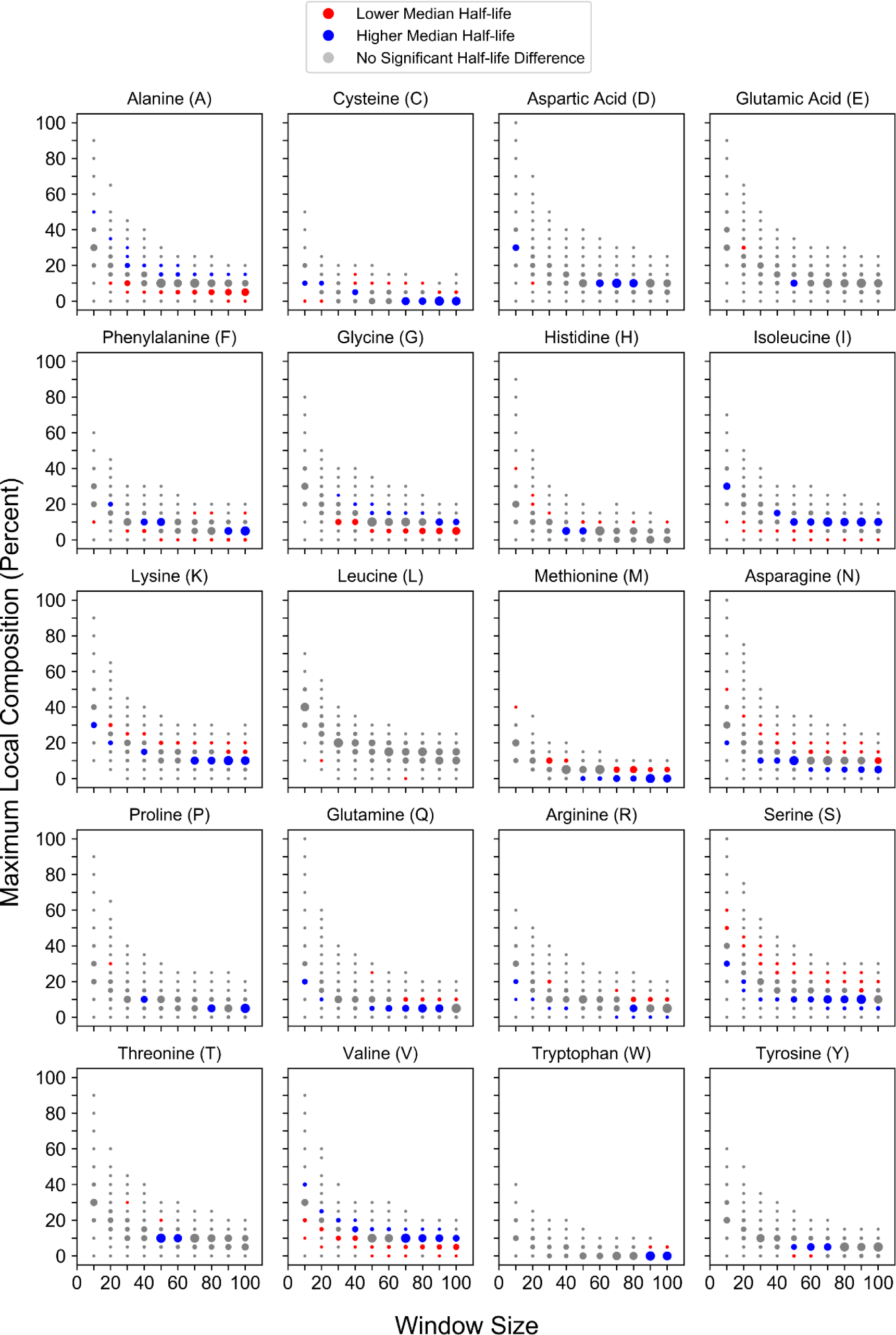
Local compositional enrichment affects the half-life of proteins lacking statistically-biased domains.

**Fig S5.**
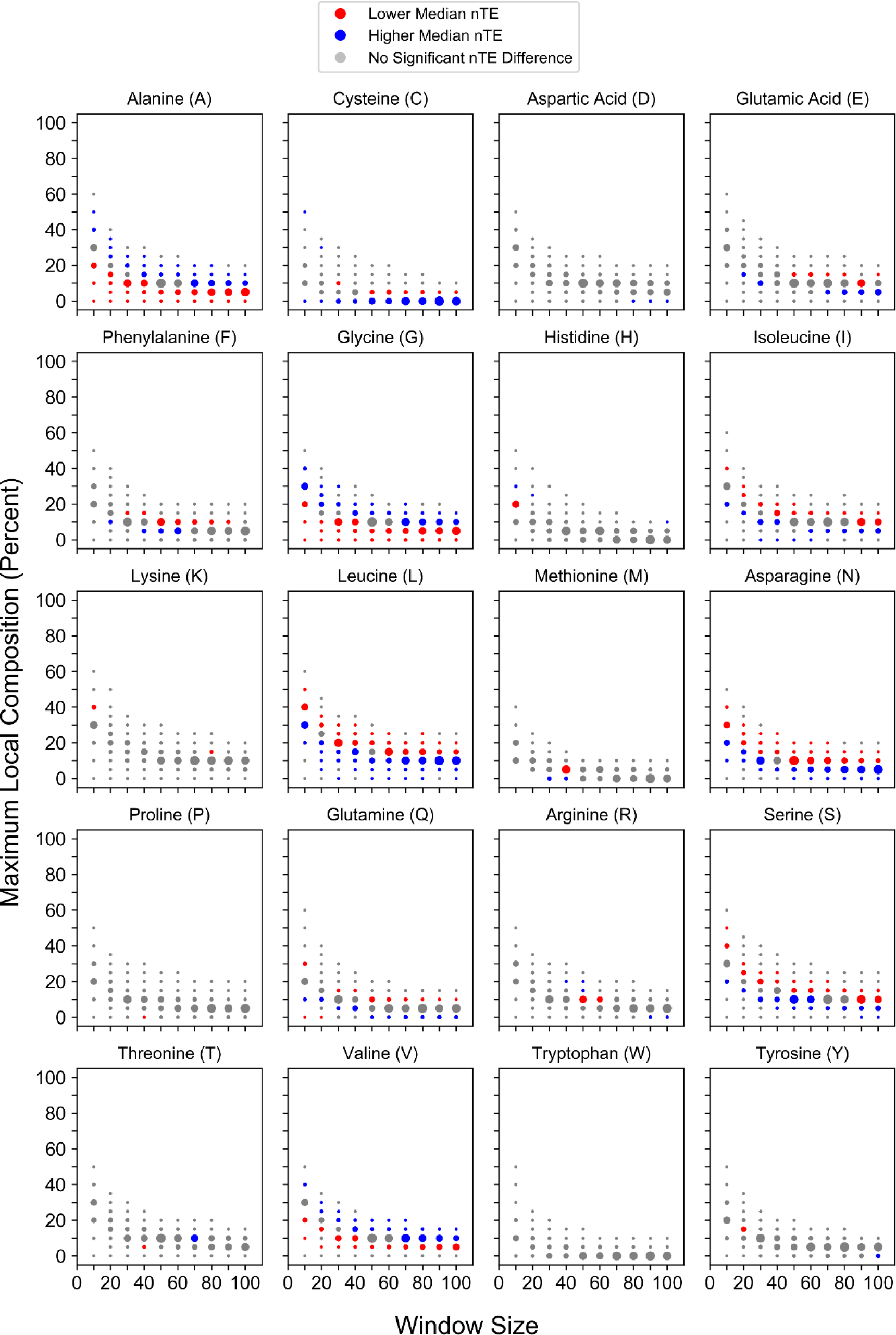
Local compositional enrichment affects the nTE of proteins lacking LCDs.

**Fig S6.**
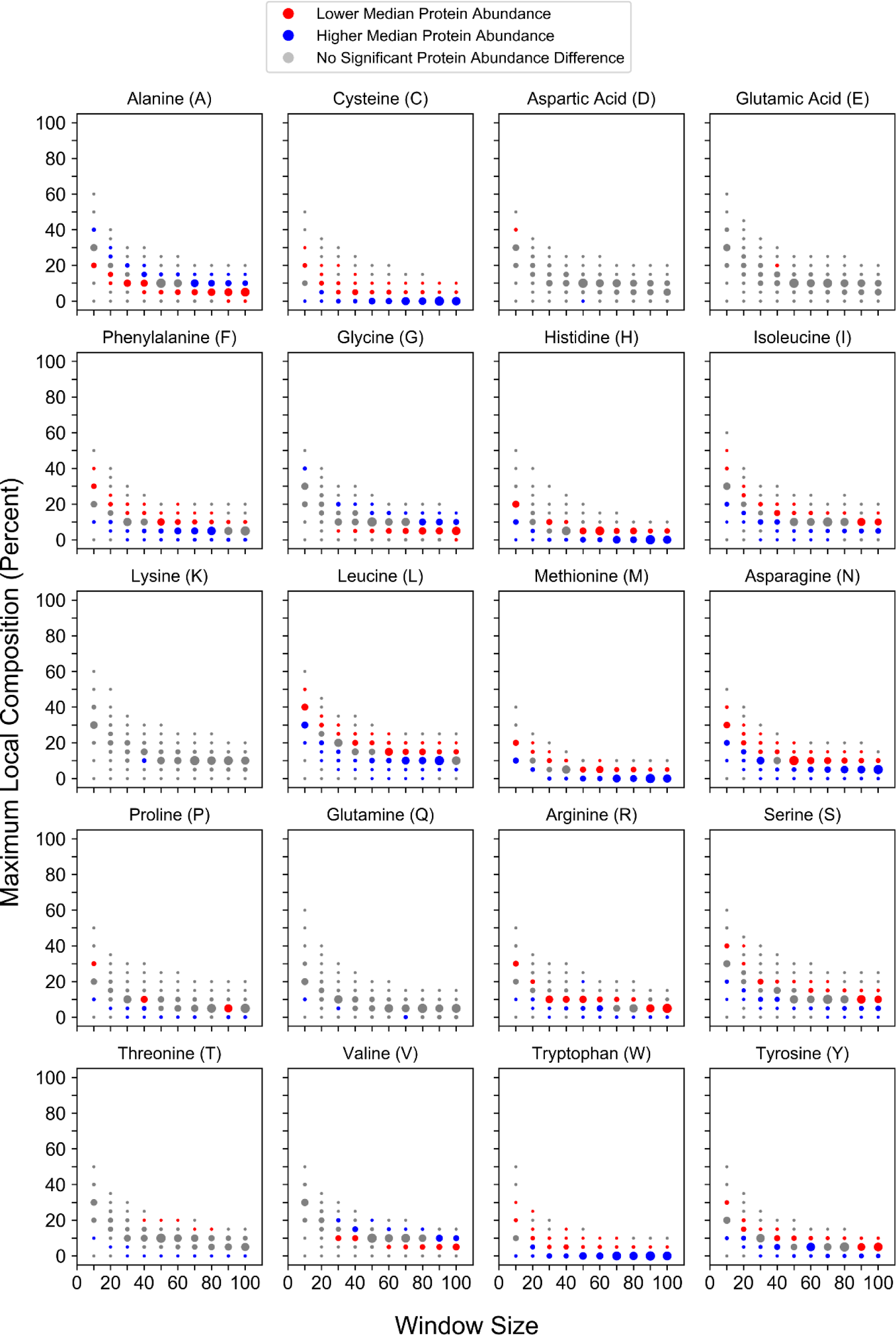
Local compositional enrichment affects the abundance of proteins lacking LCDs.

**Fig S7.**
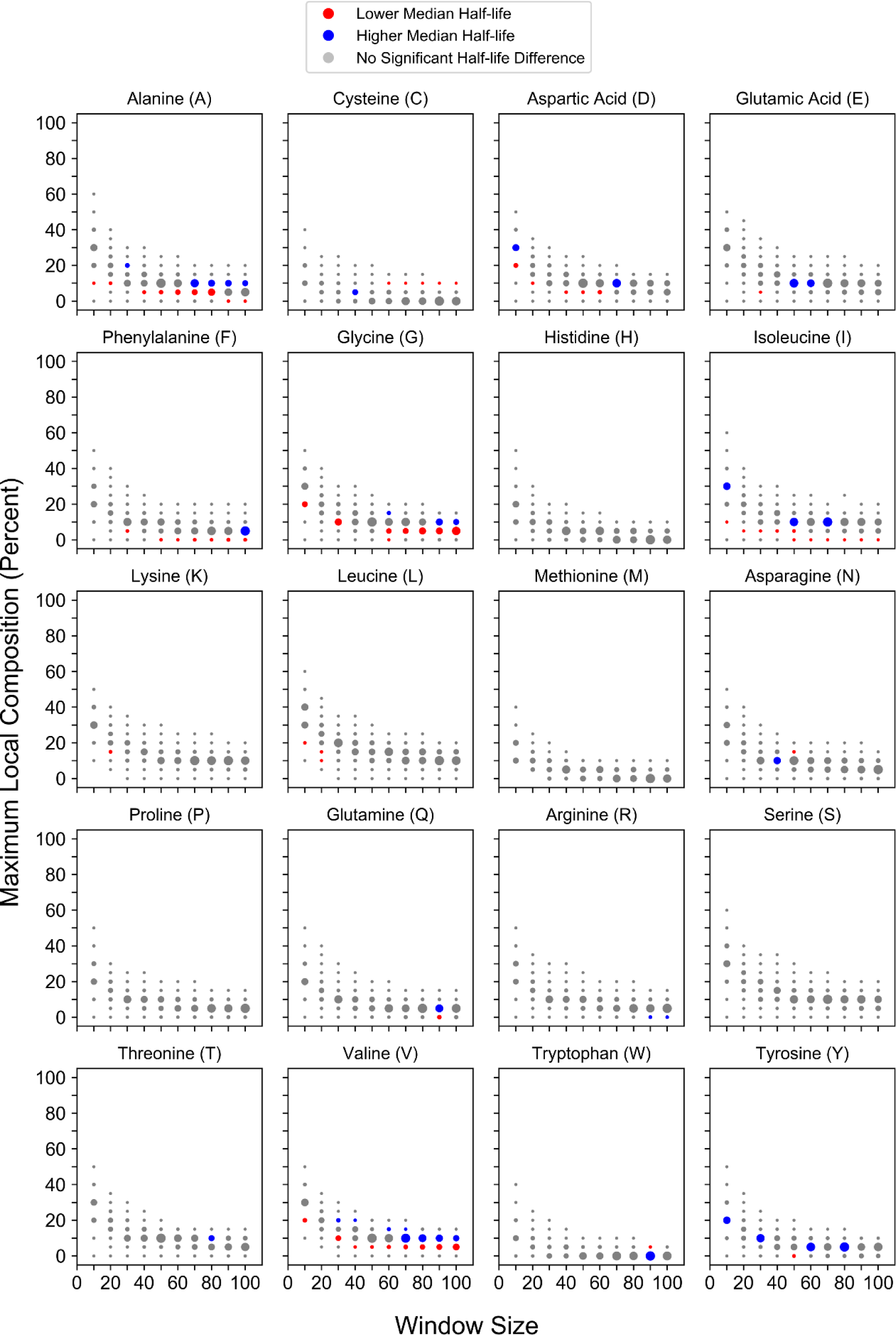
Local compositional enrichment affects the half-life of proteins lacking LCDs.

**Fig S8.**
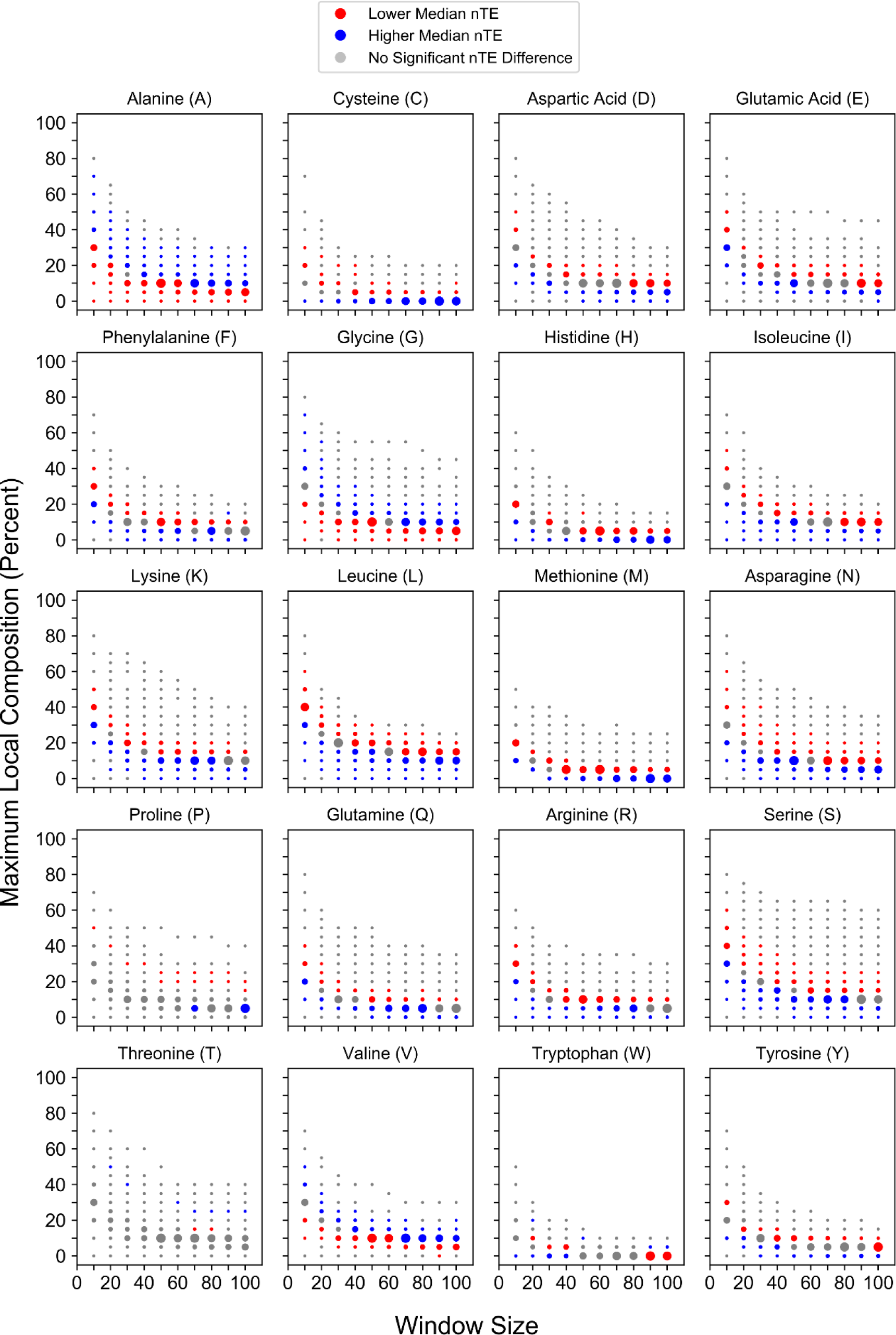
Local compositional enrichment affects the nTE of proteins lacking homopolymeric repeats.

**Fig S9.**
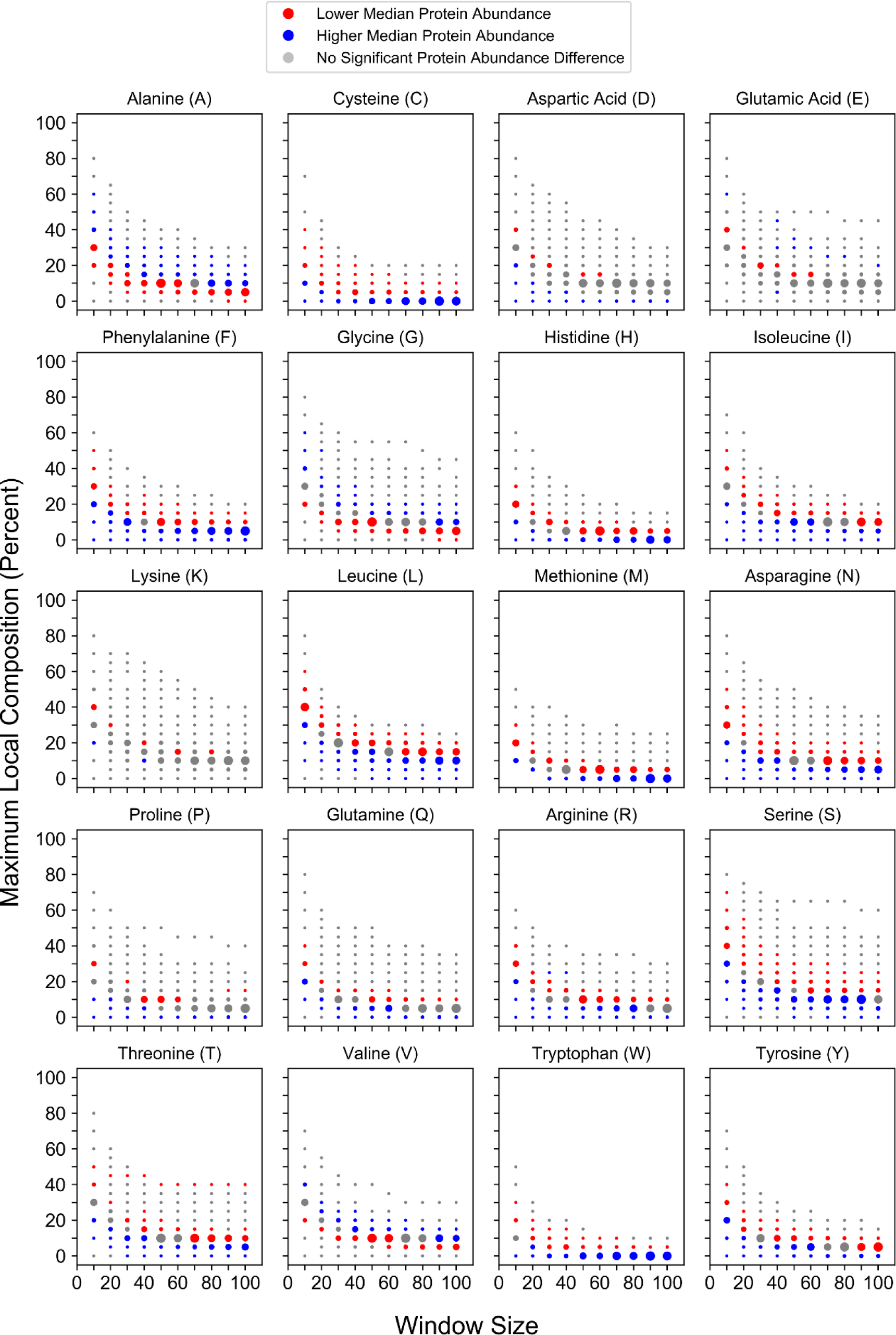
Local compositional enrichment affects the abundance of proteins lacking homopolymeric repeats.

**Fig S10.**
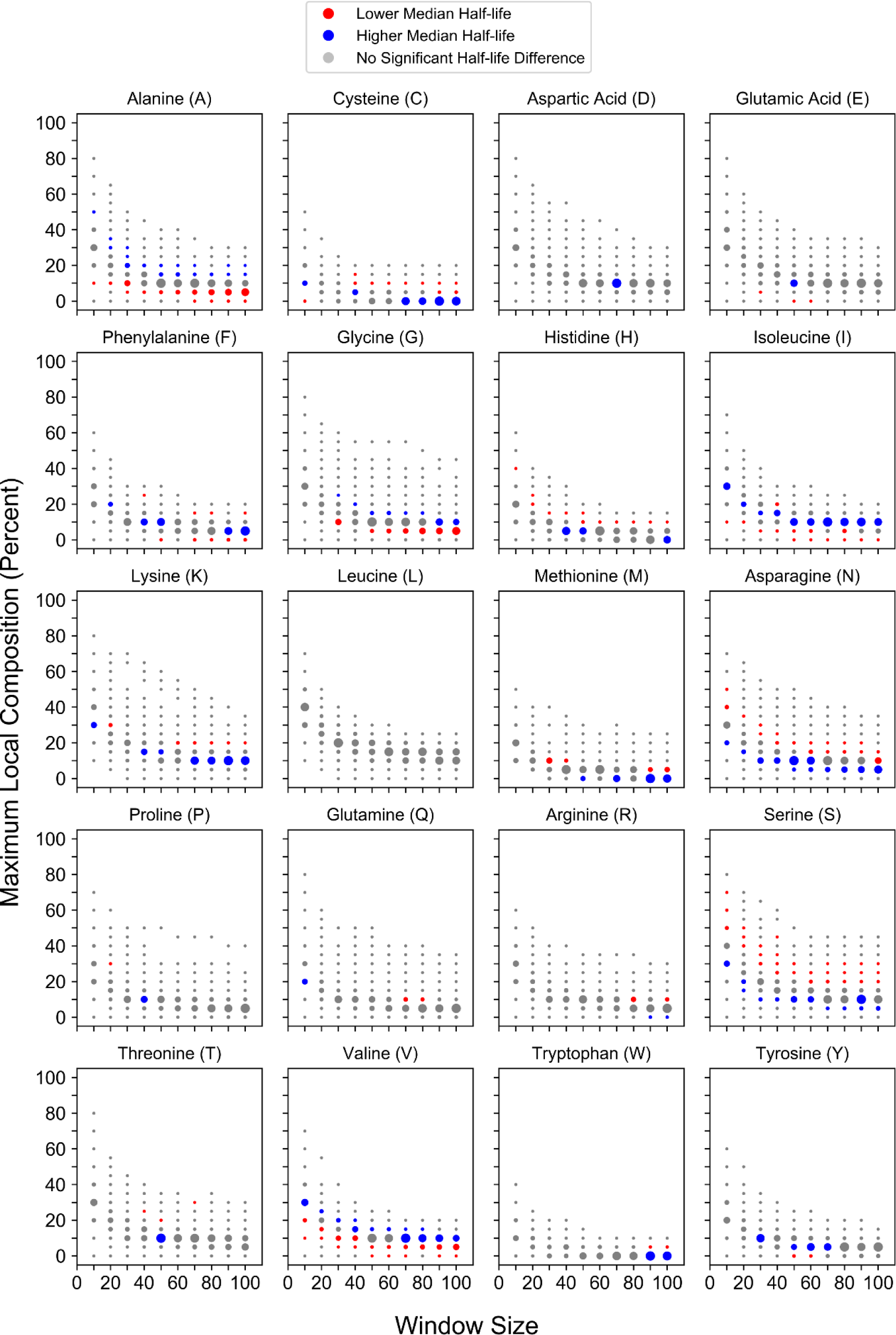
Local compositional enrichment affects the half-life of proteins lacking homopolymeric repeats.

**Fig S11.**
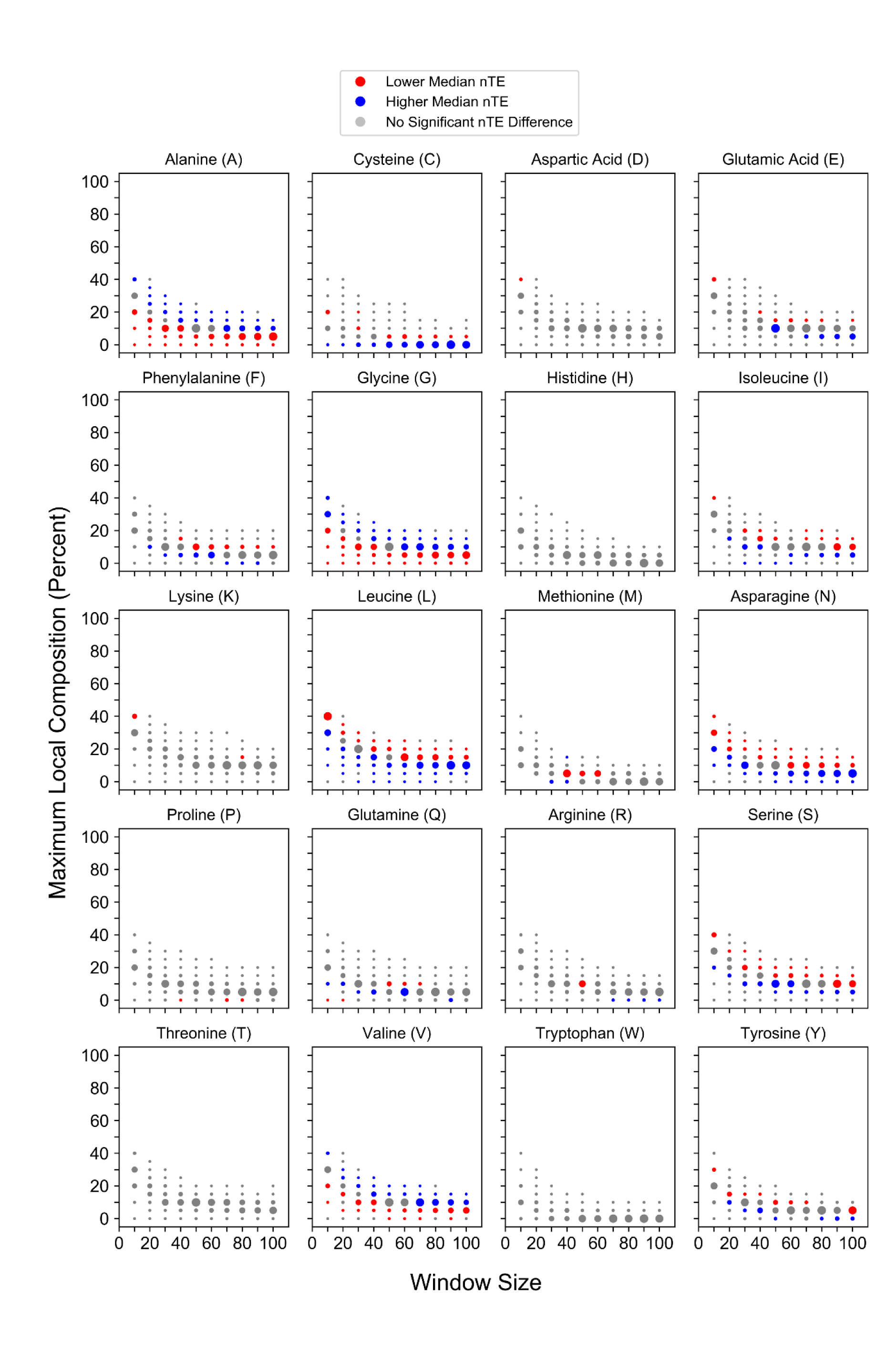
Local compositional enrichment affects the nTE of proteins lacking statistically-biased domains.

**Fig S12.**
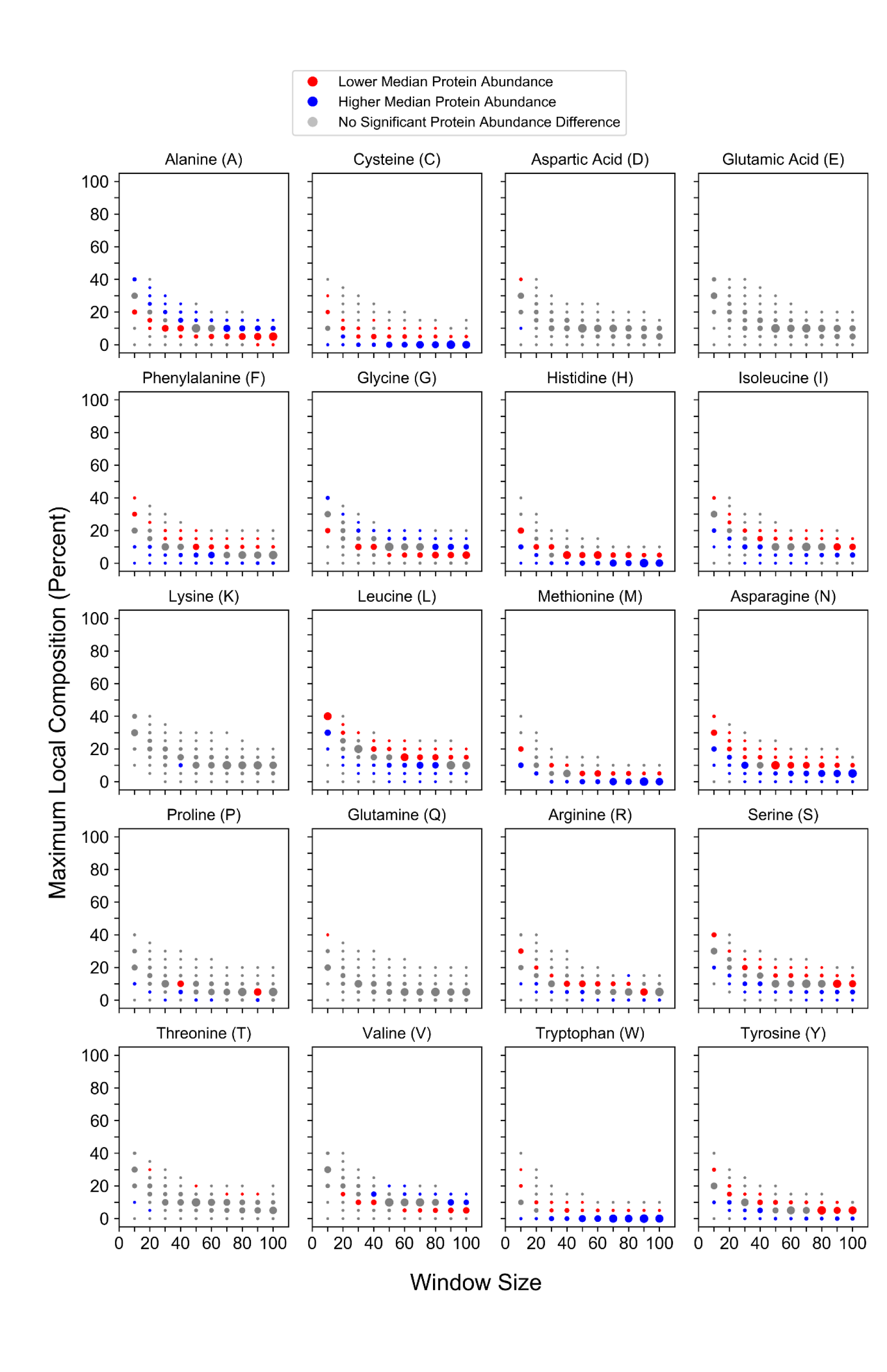
Local compositional enrichment affects the abundance of proteins lacking statistically-biased domains.

**Fig S13.**
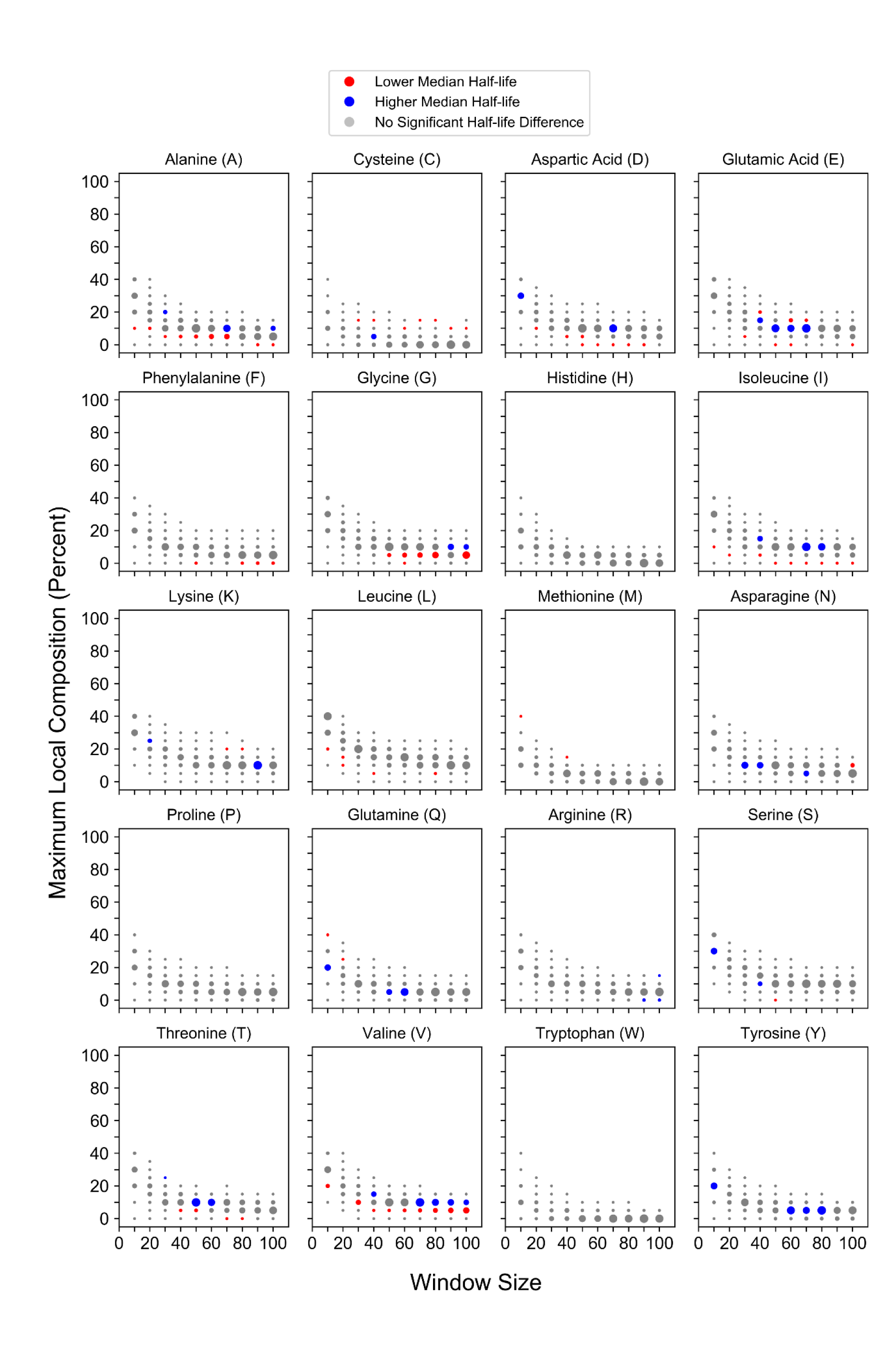
Local compositional enrichment affects the half-life of proteins lacking statistically-biased domains.

**Fig S14.**
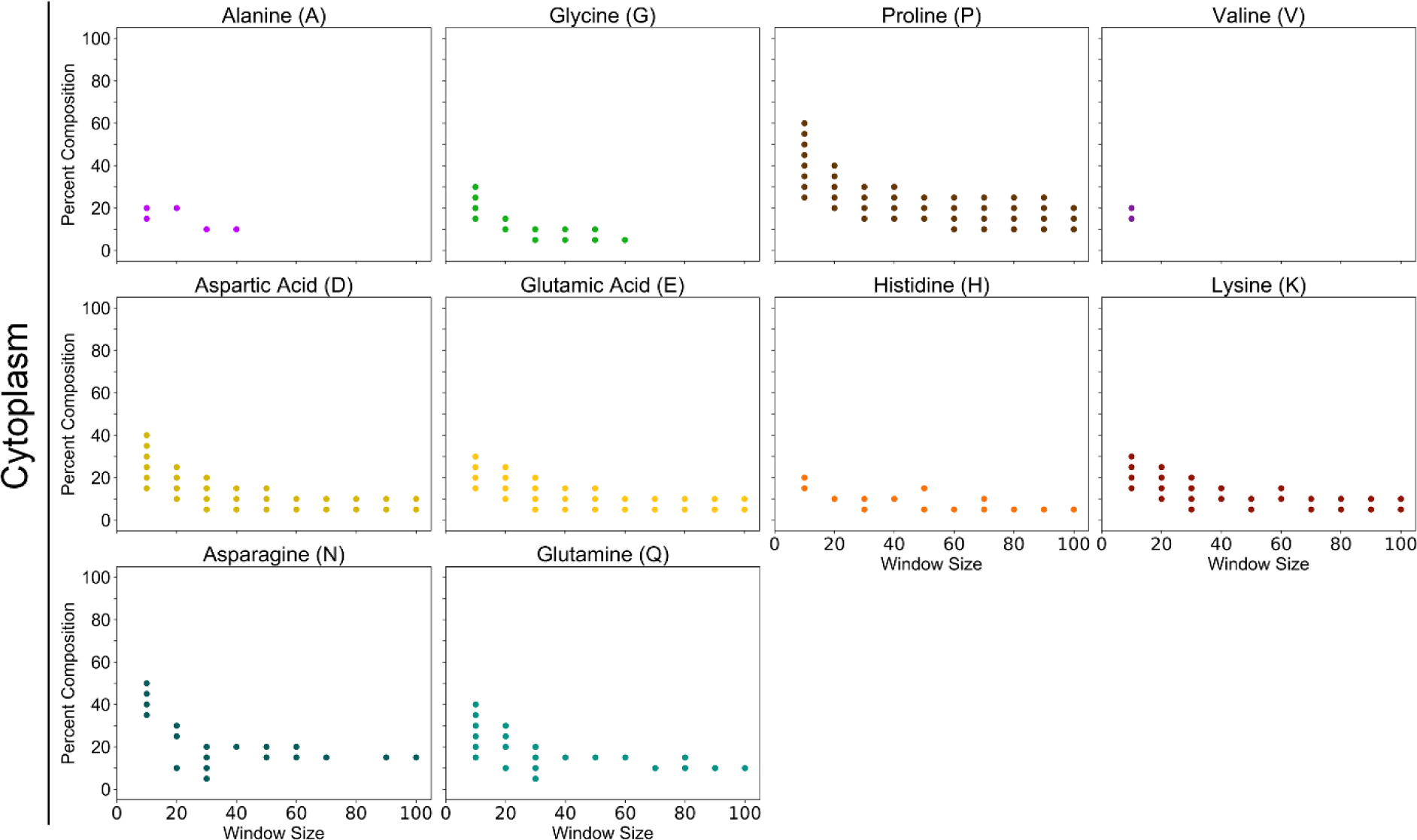
Individual amino acid composition profiles for protein sets associated with the cytoplasm. For this figure and related subsequent figures, all plotted points indicate protein sets for which association with the indicated subcellular compartment is statistically significant (Bonferonni corrected *p* < 0.05; see Methods). Plots are shown only for amino acids with at least two composition bins significantly associated with the indicated subcellular compartment.

**Fig S15.**
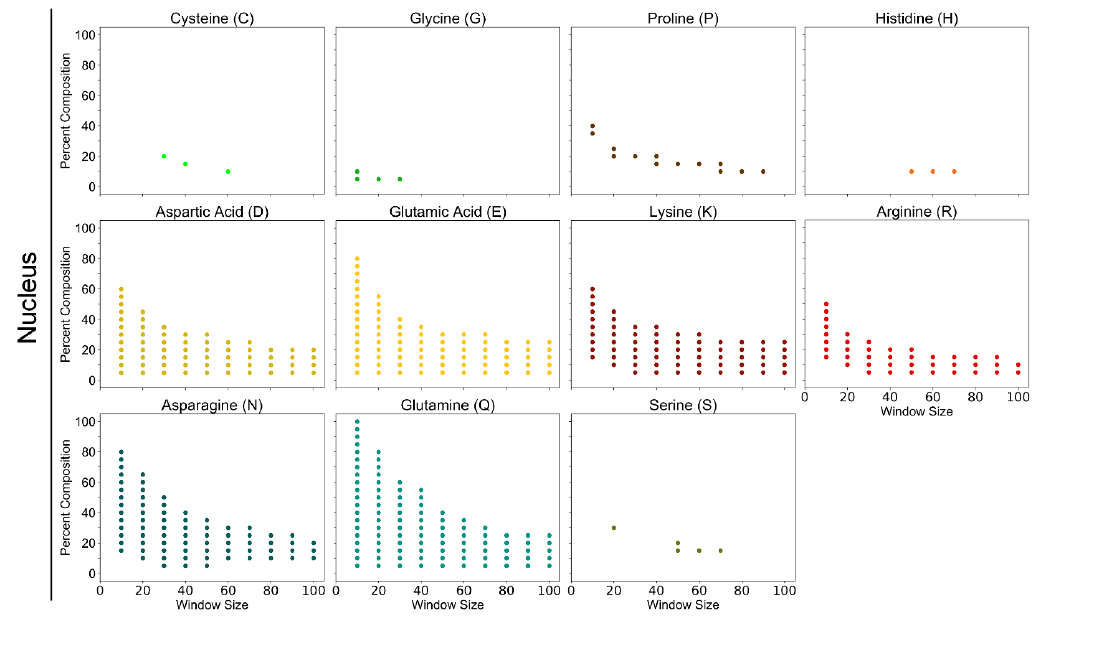
Individual amino acid composition profiles for protein sets associated with the nucleus.

**Fig S16.**
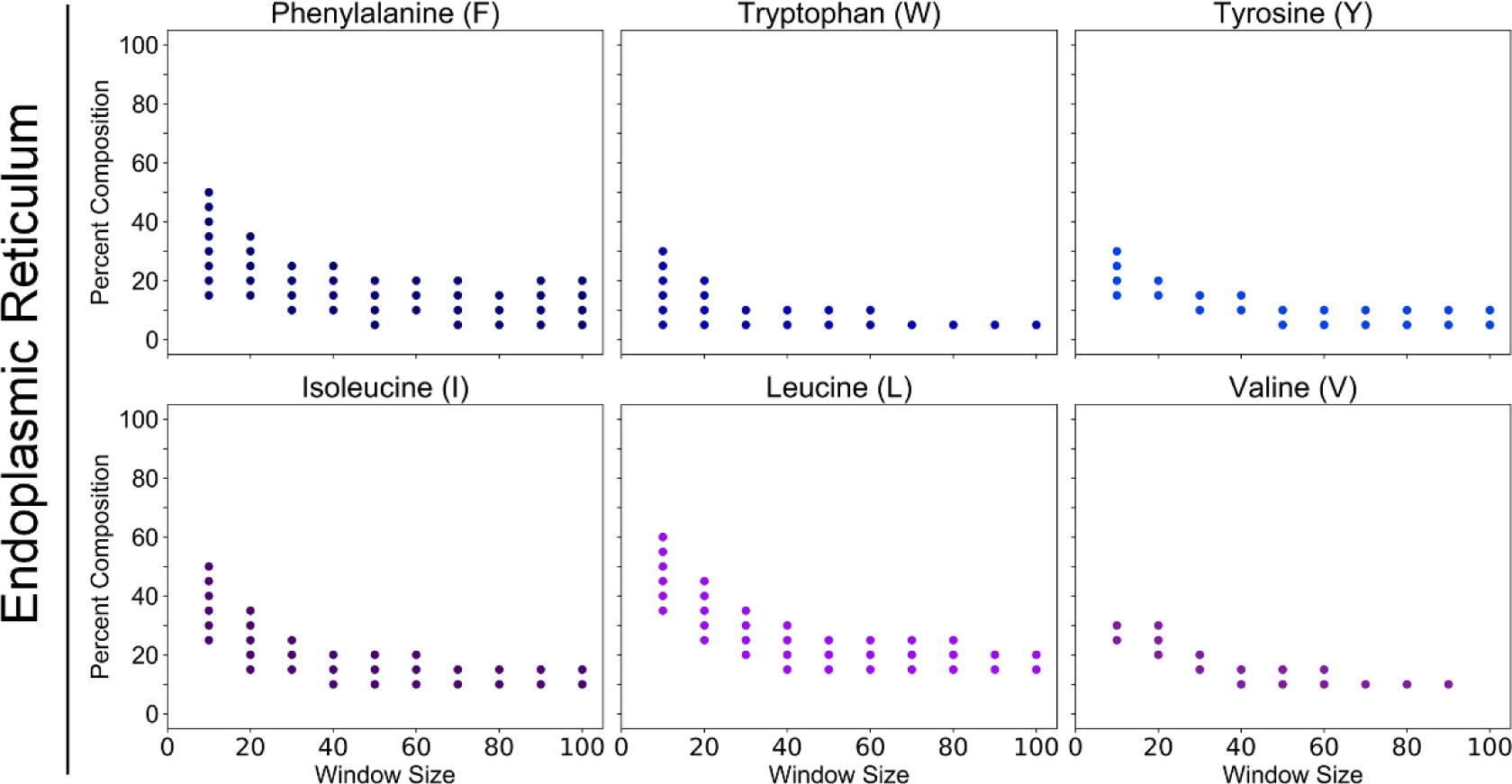
Individual amino acid composition profiles for protein sets associated with the endoplasmic reticulum.

**Fig S17.**
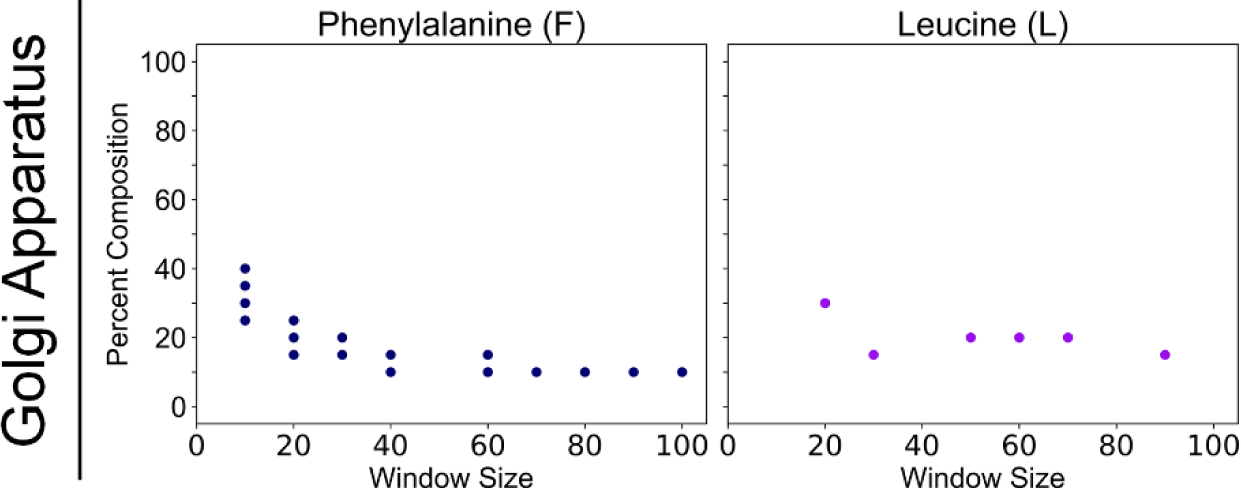
Individual amino acid composition profiles for protein sets associated with the Golgi apparatus.

**Fig S18.**
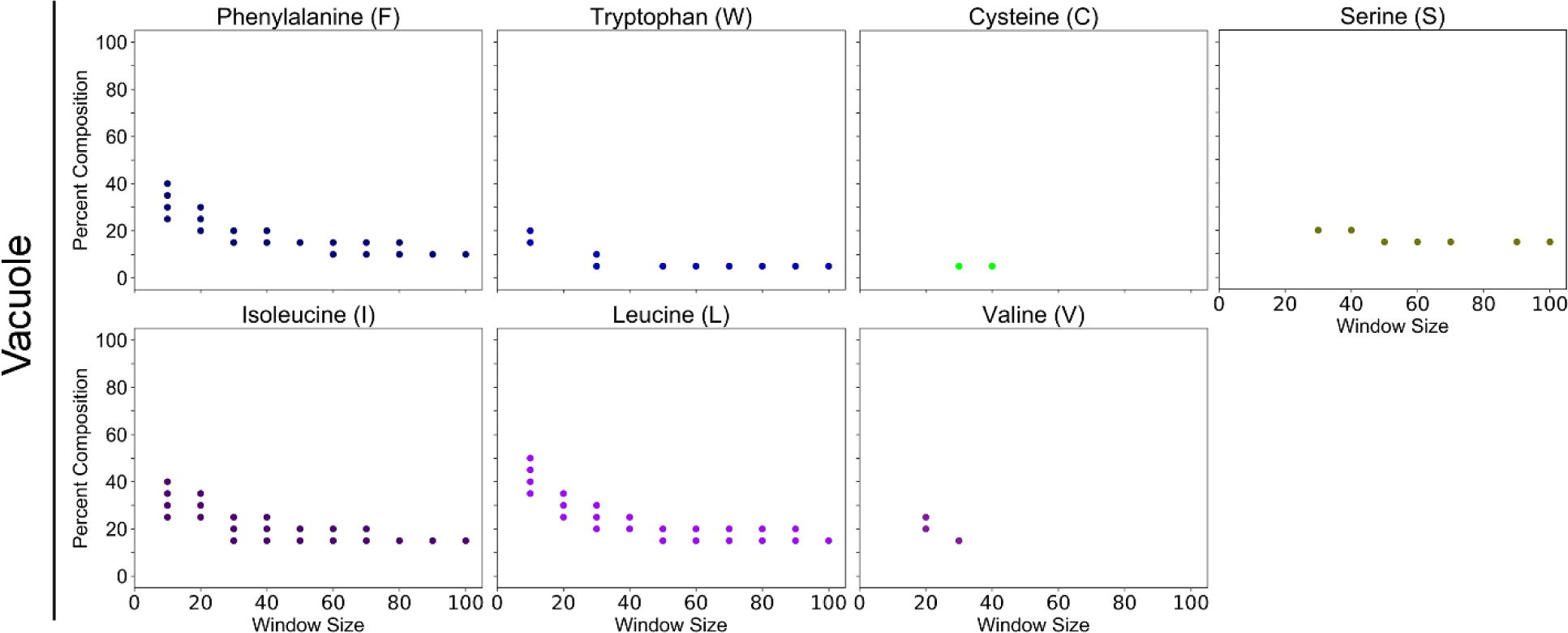
Individual amino acid composition profiles for protein sets associated with the vacuole.

**Fig S19.**
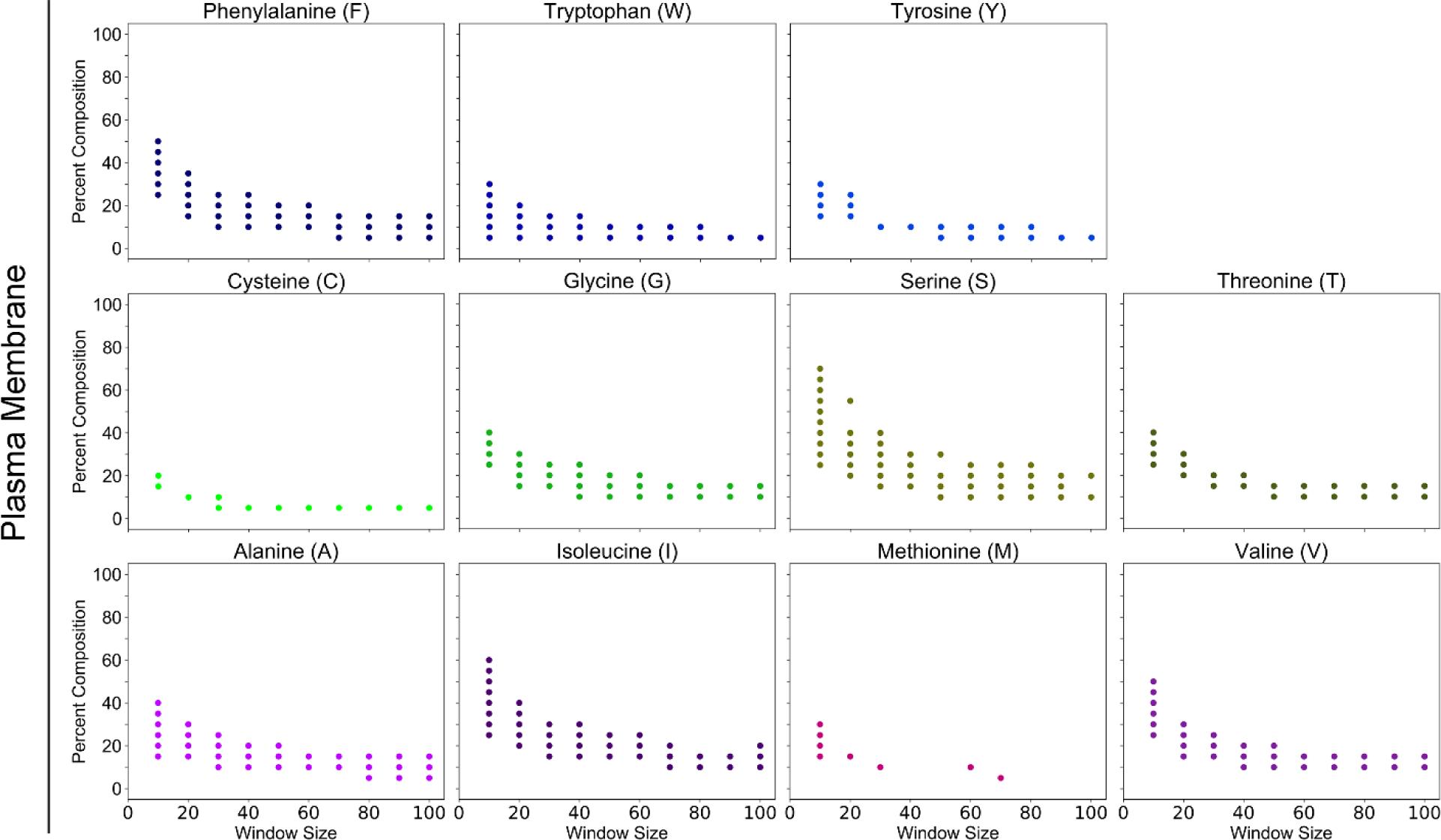
Individual amino acid composition profiles for protein sets associated with the plasma membrane.

**Fig S20.**
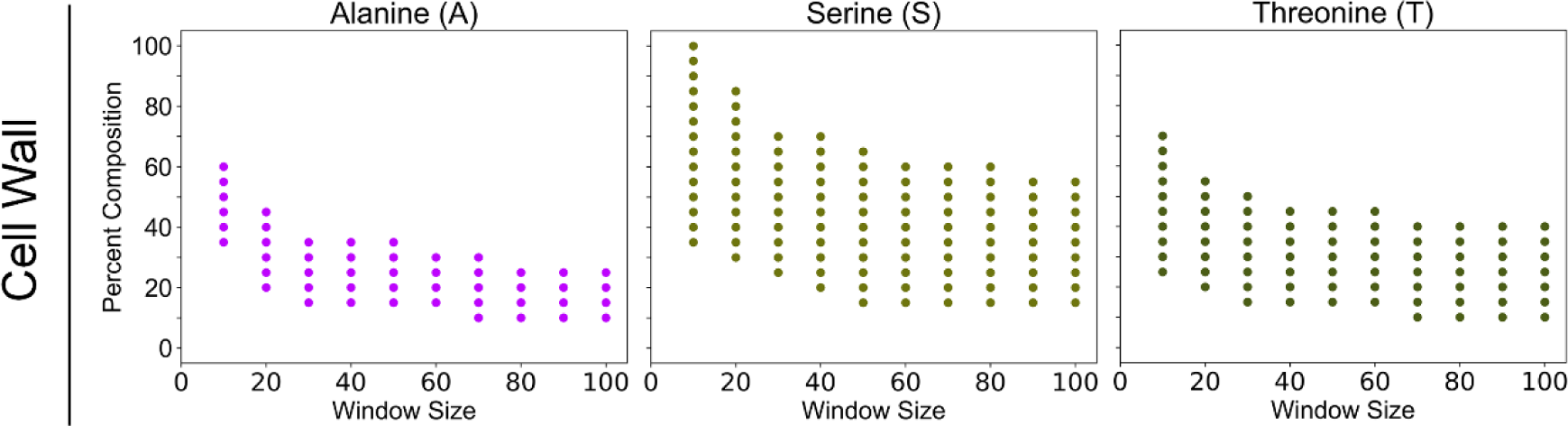
Individual amino acid composition profiles for protein sets associated with the cell wall.

**Fig S21.**
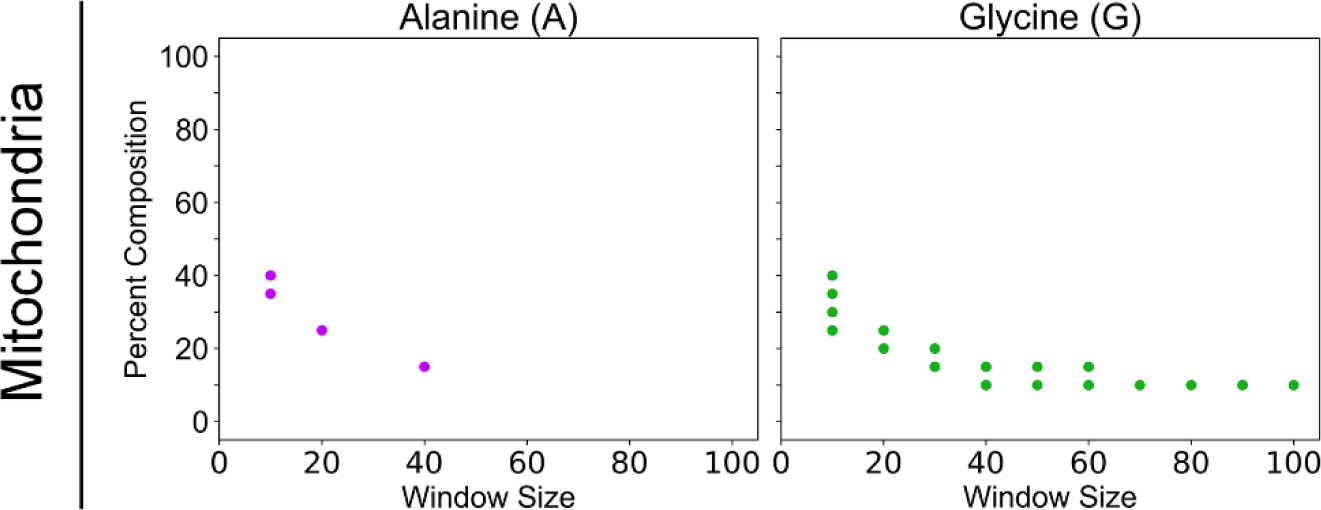
Individual amino acid composition profiles for protein sets associated with mitochondria.

**Fig S22.**
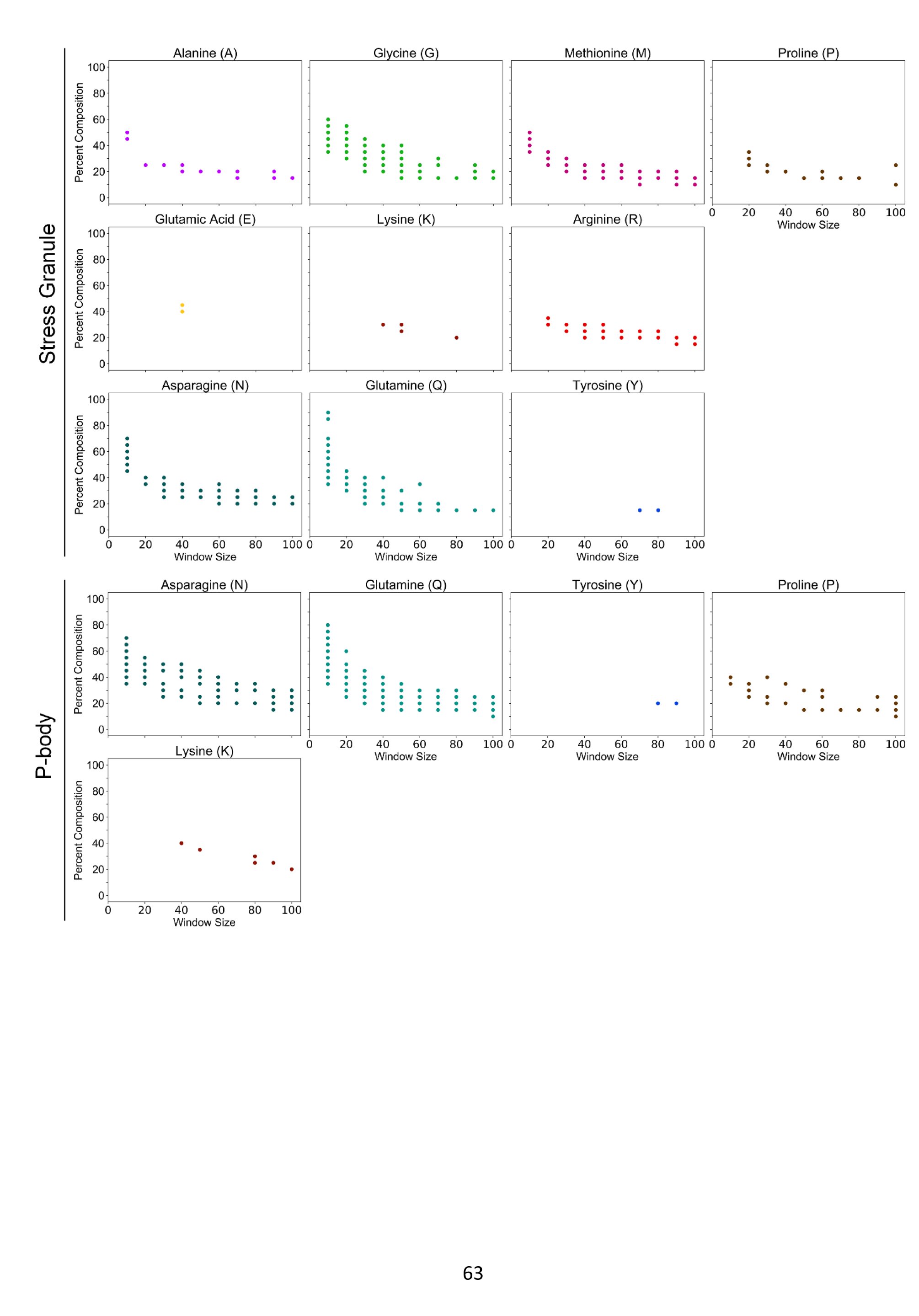

